# VSTM2L protects prostate cancer cells against ferroptosis via inhibiting VDAC1 oligomerization and maintaining mitochondria homeostasis

**DOI:** 10.1101/2024.08.03.606470

**Authors:** Juan Yang, Xiao Lu, Jing-Lan Hao, Lan Li, Yong-Tong Ruan, Xue-Ni An, Xiao-Ming Dong, Ping Gao

**Affiliations:** College of Life Sciences, Shaanxi Normal University, 710119, Xi’an, China; Institute for Research on Health Information and Technology, School of Public Health, Xi’an Medical University, 710021, Xi’an, China

## Abstract

Mitochondria play a critical role in initiating and amplifying ferroptosis. VDAC1 embedded in the mitochondrial outer membrane, exerts a crucial role in regulation of ferroptosis. However, the mechanisms of VDAC1 oligomerization in regulating ferroptosis are not well elucidated. Here, we identified that VSTM2L, a novel VDAC1 binding protein, is positively associated with prostate cancer (PCa) progression, and a key regulator of ferroptosis. Moreover, VSTM2L knockdown in PCa cells enhanced the sensibility of RSL3-induced ferroptosis. Mechanistically, VSTM2L forms complex with VDAC1 and HK2, enhancing their binding affinity and preventing VDAC1 oligomerization, thereby inhibiting ferroptosis and maintaining mitochondria homeostasis *in vitro* and *in vivo*. Collectively, our findings reveal a pivotal role for VSTM2L in driving ferroptosis resistance and highlight its potential as a ferroptosis-inducing therapeutic target for the treatment of PCa.

## INTRODUCTION

Prostate cancer (PCa) is one of the most popular cancer types threatening the health of men globally^1, 2, 3^. While radiation, surgery and androgen deprivation therapy could improve the survival rate and time of the patients, many of them eventually developed into castration resistant and metastatic prostate cancer and portends a poor prognosis^4^. At present, published evidences have indicated that ferroptosis could enhance the sensitivity of advanced PCa cells to therapeutic agents^5, 6^, exploring the key factors in regulating ferroptosis is imperative to find effective targets for PCa treatment.

Accumulated evidences indicated that mitochondria play a critical role in maintaining the biosynthetic and metabolic activities of cancer cells, and serves as key regulatory nodes in initiating and amplifying ferroptosis^7, 8, 9, 10^. Voltage-Dependent Anion Channel (VDAC), predominantly embedded in the mitochondrial outer membrane (MOM), governs mitochondrial homeostasis and distinct modes of cell death, including apoptosis, autophagy and ferroptosis^11, 12, 13, 14^. VDAC1, the most abundant among the three VDAC isoforms, has been well-documented as a crucial regulator of cell death through interactions with various proteins, including anti-apoptotic proteins (B-cell lymphoma 2 (Bcl-2), Bcl-extra large (Bcl-xL) and hexokinase (HK1 and HK2))^15, 16, 17, 18^, proapoptotic proteins (BH3-interacting domain death agonist (BID) and Bcl-2-associated X protein (Bax))^19, 20^, mitophagy-related proteins (Translocator Protein (TSPO) and Parkin)^21, 22, 23^ *et al*.. It was reported that HK2 plays an important role in keeping the monomeric status of VDAC1 in cancer cells^24^. Moreover, the oligomeric form of VDAC1, rather than its monomeric counterpart, is involved in the regulation of ferroptosis ^25, 26^. However, the regulatory mechanism underlying VDAC1 oligomerization in the modulation of ferroptosis in prostate cancer is still elusive.

In this study, we first explored the binding proteins of VDAC1 in prostate cancer cells using immunoprecipitation coupled to mass spectrometry (IP/MS) analysis, and found a novel interacting protein of VDAC1, V-Set and Transmembrane Domain Containing 2 Like (VSTM2L). VSTM2L, also known as C20orf102, was initially found to interact with Humanin (HN), and was selectively expressed in the central nervous system. VSTM2L is a secreted antagonist of HN and could play a role in regulating HN biological functions^27^. Recent studies suggested that VSTM2L is recognized as a potential biomarker closely associated with cancer cell metastasis and prognosis^28, 29^. Thus far, the biological roles and the underlying molecular mechanisms of VSTM2L in PCa cells are still blurry.

Here, our study showed that the VDAC1 binding protein VSTM2L was a novel oncogene and ferroptosis suppressor in PCa cells. VSTM2L knockdown impeded cell growth and migration by facilitating ferroptosis of PCa. Molecular biology studies uncovered that VSTM2L is a crucial factor to keep the interaction between VDAC1 and HK2. Decreased VSTM2L led to the dissociation of HK2 from VDAC1 and the increased oligomerization of VDAC1, which disturbed the homeostasis of mitochondria and triggered the ferroptosis of PCa cells. Our findings not only revealed a novel link between VSTM2L and ferroptosis but also disclosed the regulatory mechanism underlying VDAC1 oligomerization in the modulation of ferroptosis, suggesting that VSTM2L is a predictive factor with therapeutic potential for PCa treatment.

## RESULTS

### VSTM2L is a novel VDAC1 binding protein in PCa Cells

Considering the crucial role of VDAC1 in maintaining the homeostasis of mitochondria, we firstly explored the interacted proteins with VDAC1 in PCa cells by IP-MS analysis. From the top five scored proteins, we noticed that VSTM2L is the newly identified VDAC1-interacting protein and the functional role of VSTM2L in PCa cells has never been reported (Fig. 1A, Fig. S1) (Supplementary File 1). To confirm, co-immunoprecipitation (co-IP) assay was performed in VDAC1 and VSTM2L co-transfected HEK293T cells, the results showed VDAC1 interacted with VSTM2L (Fig. 1B-C). Moreover, the semi-endogenous interaction between VSTM2L and VDAC1 was determined in Myc-VDAC1 overexpressed DU145 or 22Rv1 cell lines by Co-IP assay (Fig. 1D-E). To further evaluate whether there is direct interaction between VSTM2L and VDAC1, we performed the Glutathione S-transferase (GST) pull-down assay and confirmed the direct binding of VSTM2L to VDAC1 (Fig. 1F). In addition, Coomassie Brilliant Blue staining of the IP cell lysates revealed that VSTM2L was successfully pulled down and further to LC-MS (Fig. 1G). Venn diagram revealed that 574 proteins were overlapped in VSTM2L and VDAC1 binding proteins (Fig. 1H) (Supplementary File 2). Kyoto Encyclopedia of Genes and Genomes (KEGG) pathways analysis of these 574 proteins further revealed that these proteins were highly enriched in Citrate cycle (TCA cycle), Glycolysis and Fatty acid metabolism pathways (Fig. 1I). These results indicated that VSTM2L might play an important role in regulating mitochondria metabolism. In order to clarify the main localization of VSTM2L in the PCa cells, We have separately extracted the mitochondria and cytoplasmic proteins from three prostate cancer cell lines (PC3, DU145, 22Rv1), respectively, and tested the protein levels of VSTM2L. As shown in Figure 1J, VSTM2L indeed mainly located in the mitochondria rather than in the cytoplasm in PC3, Du145 and 22Rv1 cells. Moreover, Immunofluorescence (IF) staining showed that endogenous VSTM2L was localized to mitochondria in Du145 cells (Fig. 1K).

**Fig. 1.**
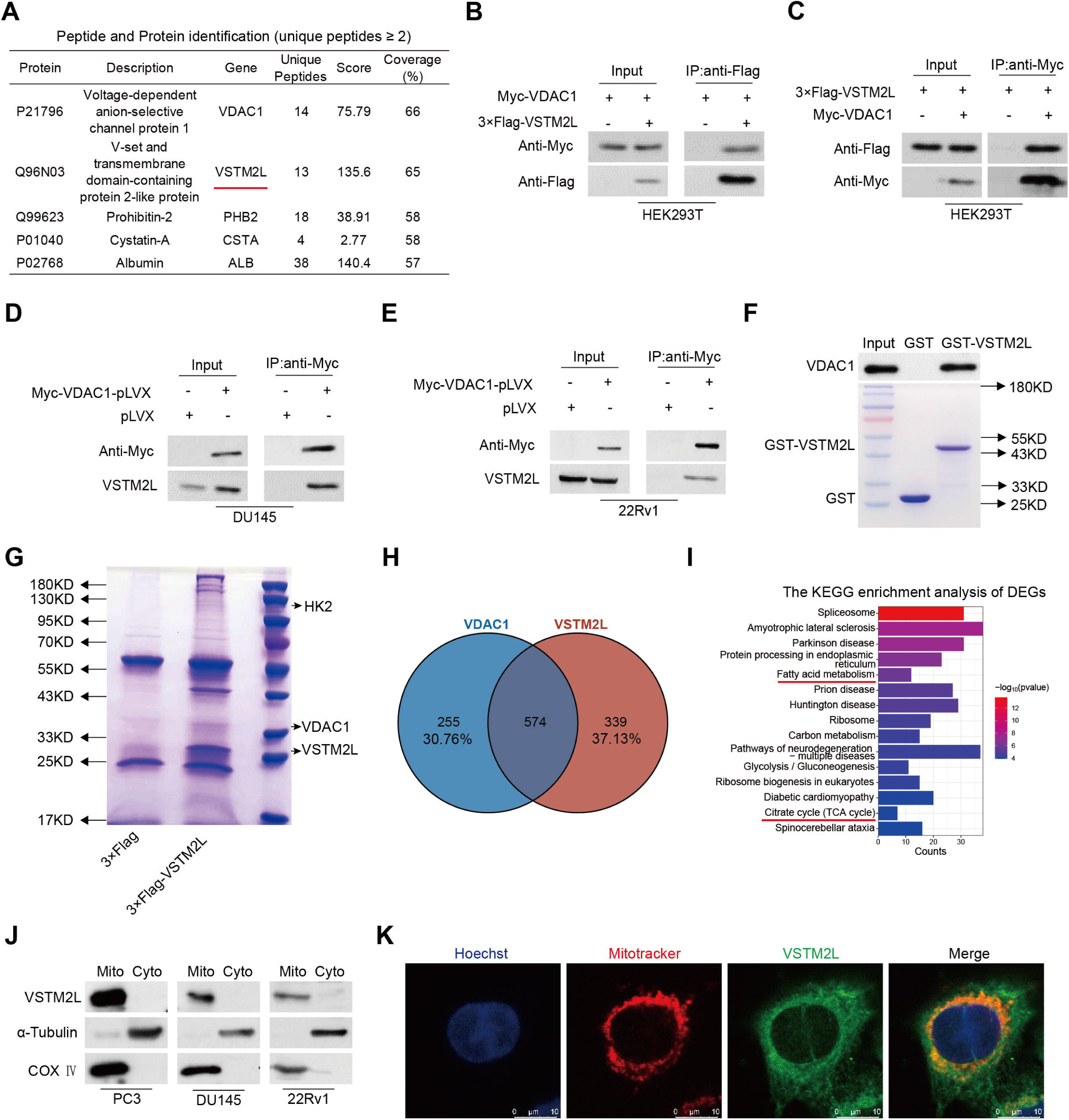
VSTM2L is a novel VDAC1 interacting protein in Pca cells. **A** The top five proteins, identified by co-IP/MS, was elucidated in association with VDAC1 in LNCaP cells. **B, C** HEK293T cells were transfected with Myc-VDAC1 and 3×Flag-VSTM2L expression plasmids as indicated, whole cell lysates were extracted and immunoprecipitated with anti-Flag (**B**) or anti-Myc (**C**) antibodies. **D, E** DU145 (**D**) and 22Rv1 (**E**) cells overexpressed Myc-VDAC1-pLVX or control cells were subjected to immunoprecipitation with anti-Myc antibody and followed by western blot analysis. **F** The direct interaction between VSTM2L and VDAC1 was confirmed by GST pull-down assay. GST-VSTM2L fusion proteins were purified and incubated with PC3 cell lysates and analyzed by western blotting. Purified GST proteins served as negative controls. **G** A representative image of an SDS-PAGE gel stained with Coomassie blue. LNCaP cells transfected with 3×Flag-VSTM2L were collected for immunoprecipitation with anti-Flag antibodies, and separated by SDS-PAGE gel. The gel was subsequently stained with Coomassie blue to visualize the distinct binding bands. **H** Venn diagram showing the 574 genes obtained by intersecting the IP-MS data of VDAC1 and VSTM2L. **I** KEGG pathways enriched for the 574 genes obtained. **J** Western blotting analysis of the expression of VSTM2L in mitochondria and cytoplasm of PC3, DU145 and 22Rv1 cells, respectively. Mitochondria and cytoplasmic protein levels were normalized to COX Ⅳ and α-Tubulin, respectively. **K** Representative images of immunofluorescence staining for VSTM2L and MitoTracker in DU145 cells. The nucleus was stained with Hoechst. Scale bars, 50 μm and 10 μm.

Taken together, these results declared that VSTM2L is a novel binding protein of VDAC1 in PCa cells, and VSTM2L with mitochondrial localization might play a pivotal role in regulating mitochondria metabolism.

### VSTM2L is positively associated with prostate cancer aggressiveness

To investigate the functional role of VSTM2L in prostate cancer, we first analyzed the top 25 over-expression genes in prostate adenocarcinoma (PRAD) using the published cohort from TCGA. As shown in Fig. 2A, VSTM2L ranked as the 24th over-expression gene in PRAD. Further, we observed a marked elevation of VSTM2L mRNA levels in tumor specimens as compared to normal controls based on the GEPIA, GENT2 and UALCAN prostate patient cohorts, respectively (Fig. 2B-C; Fig. S2 A). Then, we measured the mRNA levels of VSTM2L in different PCa cell lines, and observed that VSTM2L was highly expressed in multiple PCa cell lines (22Rv1, DU145 and PC3) in compare with normal prostate epithelial RWPE-1 cells (Fig. S2 B). Moreover, we collected 78 pairs of prostate tumor and adjacent tissues and examined the protein levels of VSTM2L using immunohistochemistry (IHC) staining assay. The results indicated that the expression levels of VSTM2L were up-regulated in the tumor tissues compared to the adjacent tissues (Fig. 2D-E). In addition, a higher expression of VSTM2L referred to a poor prognosis of PRAD patients according to the Kaplan-Meier progression free survival (PFS) and disease-free survival (DFS) analysis with the cohort from cBioPortal (Fig. 2F-G; Fig. S2 C-E). Overall, these results suggested that the expression levels of VSTM2L were apparently elevated in prostate cancer samples and associated with shorter survival of patients.

**Fig. 2.**
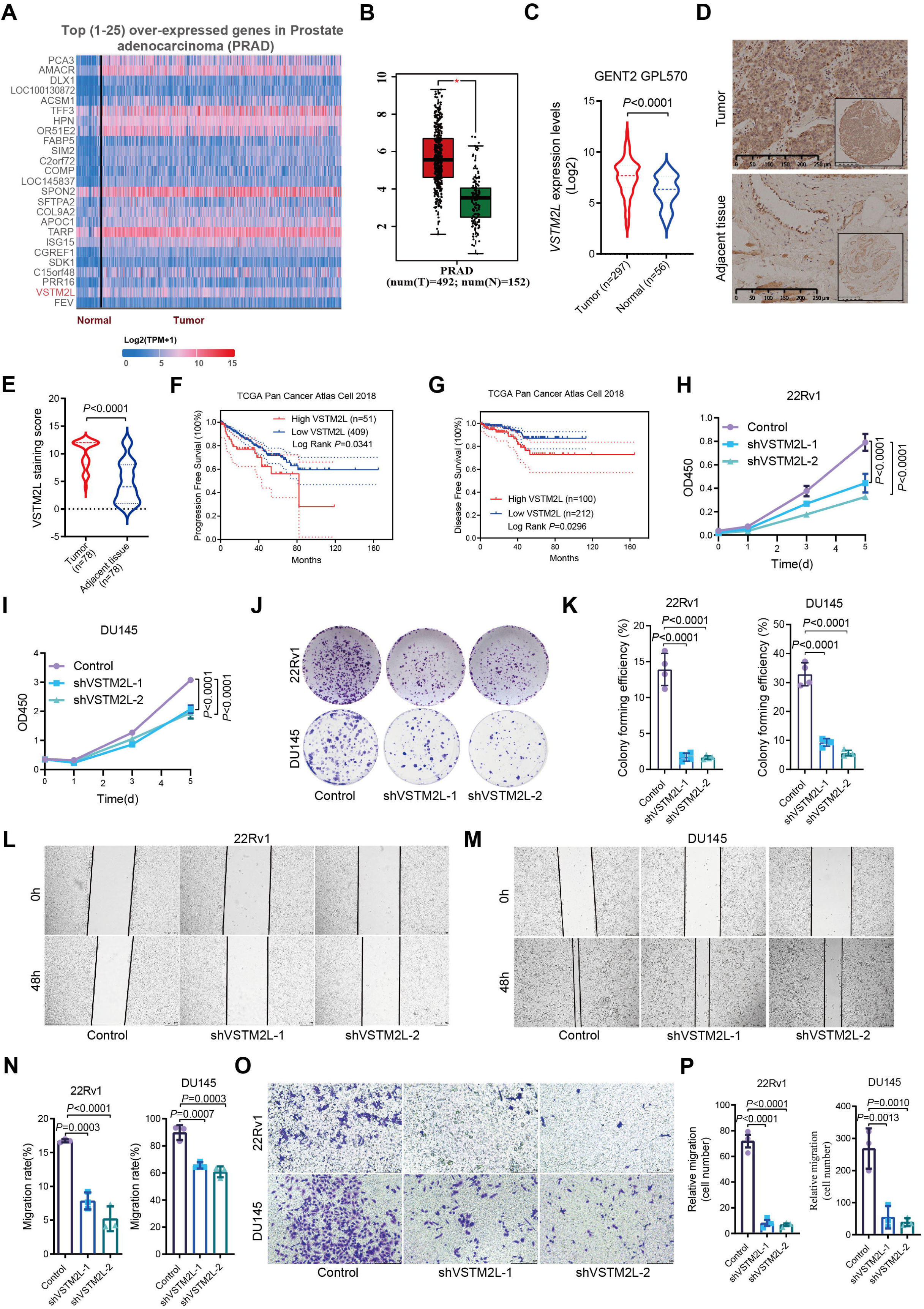
VSTM2L is positively associated with prostate cancer aggressiveness. **A** The top (1-25) over-expressed genes in Prostate adenocarcinoma (PRAD) were analyzed based on the TCGA cohort by using the UALCAN database. **B-C** The mRNA levels of *VSTM2L* in PRAD and normal tissues from GEPIA (**B**) and GENT2 (**C**) database. * *P*<0.05. **D, E** The protein levels of VSTM2L from 78 paired clinical prostate cancer specimens. The IHC staining score was used to quantify the protein levels of VSTM2L. Scale bar =250 μm. **F, G** Association between Progression-free-survival (**F**) or Disease-free survival (**G**) of prostate cancer patients and *VSTM2L* mRNA expression from the TCGA database. **H, I** The cell proliferation was measured in VSTM2L knockdown 22Rv1 (**H**) or DU145 (**I**) cells by CCK8 assay. Data are shown as the mean ± SD of triplicate independent sets of experiments. **J, K** The effects of VSTM2L knockdown on the growth of 22Rv1 or DU145 cells, as detected using the colony formation assay. Data are presented as mean ± SD, and all data represent the results of three independent experiments. **L-N** Wound healing analysis for assessing migration of the indicated cell strains at 0 h and 48 h. Representative images (**L, M**) and quantification (**N**) are shown as indicated. Data from independent experiments are presented as the mean ± SD, n=3 randomly selected magnification fields, Scale bar =250 μm. **O, P** Representative pictures (**O**) and quantification analysis (**P**) of migration assays in control and VSTM2L knockdown 22Rv1 cells or DU145 cells. Data are plotted as mean ± SD, n=3 randomly selected magnification fields, Scale bar =250 μm. *P* value was determined by unpaired two-tailed Student’s *t*-test (**B, C**), paired two-tailed Student’s *t*-test (**E**), log-rank test (**F, G**), two-way ANOVA (**H, I**) and one-way ANOVA (**K, N, P**). * *P*<0.05.

We next explore the biological function of VSTM2L in PCa cells. Prostate cancer cell lines (22Rv1, DU145 and PC3 cells) were infected with lentivirus-based VSTM2L shRNA or scramble shRNA (Control) to obtain the stable strains of VSTM2L knockdown. The knockdown efficiency was detected by real-time PCR and western blot. As shown in Fig. S2 F-G, the VSTM2L shRNA lentivirus significantly decreased the mRNA and protein levels of endogenous VSTM2L compared with the control group. Then, the effect of VSTM2L knockdown on cell proliferation was investigated by CCK-8 assay. The results showed that knockdown of VSTM2L in PCa cell lines (22Rv1, DU145 and PC3 cells) significantly decreased cell proliferation ability compared with control group (Fig. 2H-I; Fig. S2 H). Suppression of VSTM2L expression in PCa cell lines also weakened cell colony formation ability (Fig. 2J-K; Fig. S2 I). In addition, inhibition of VSTM2L expression restrained the migration abilities of PCa cell lines such as 22Rv1, DU145 and PC3 cells (Fig. 2L-P; Fig. S2 J-M). Taken together, these results indicated that VSTM2L deletion inhibited cell growth and migration in PCa cells, and VSTM2L might be a novel therapeutic target for PCa treatment in clinical.

### VSTM2L suppression promotes ferroptosis of PCa Cells

After diminished the expression of VSTM2L in 22Rv1 or Du145 cells, we observed the cells with less cellular antenna and the shape of the cells turned round (Fig. 3A). This phenotype suggested that the subcellular structure of VSTM2L inhibited 22Rv1 or Du145 cells might have changed. To investigate the impact of VSTM2L on the subcellular structure of PCa cells, the transmission electron microscopy assay was performed after VSTM2L knockdown in 22Rv1 cells. At the cellular level, we observed a remarkable shrinkage of the cells with shRNA targeting VSTM2L in compare with control cells (Fig. S3 A). At the subcellular level, we observed remarkable alterations of the morphology of mitochondria in VSTM2L knocked down 22Rv1 cells, mainly characterized by a general shrink in size, increase in membrane density, and diminution in cristae (Fig. 3B). In addition, living mitochondria imaging showed a reduction in mitochondria number, mean mitochondria area, mean mitochondria perimeter, mean mitochondria form factor and mean mitochondria aspect ratio in VSTM2L inhibited DU145 and 22Rv1 cells compared with control group. (Fig. 3C-E; Fig. S3 B). Furthermore, the network of mitochondria significantly changed in these cells as well (Fig. 3F; Fig. S3 C). These alterations induced by suppression of VSTM2L expression in prostate cancer cells closely resemble the morphological characteristics observed in cells undergoing ferroptosis^30^.

**Fig. 3.**
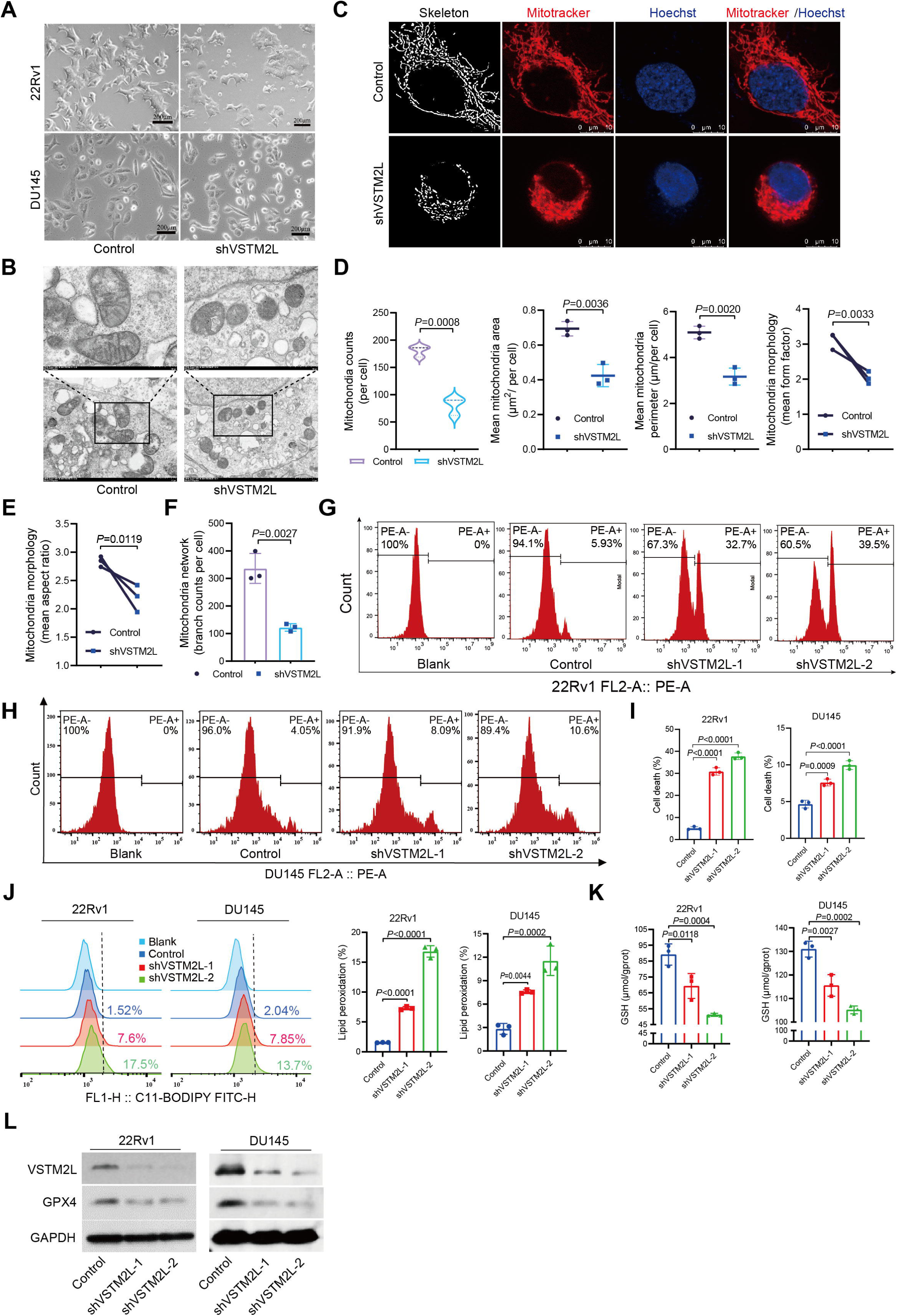
VSTM2L suppression promotes ferroptosis of PCa Cells. **A** The cell morphology of 22Rv1 and Du145 cells with VSTM2L shRNA (shVSTM2L) or control shRNA (control) were observed by inverted light microscopy. Scale bars = 200μm. **B** Representative transmission electron microscopy (TEM) images of 22Rv1 cells with control or shVSTM2L. Scale bars = 1 μm / 500 nm. **C-F** Representative images (**C**) and quantification analysis (**D-F**) of high resolution laser confocal microscopy for the morphology of mitochondria in DU145 cells transfected with control or shVSTM2L. Data shown as mean ± SD of n=3 technical replicates, Scale bars = 10 μm. **G-I** The 22Rv1(**G**) and DU145 (**H**) cells with VSTM2L shRNA (shVSTM2L-1, shVSTM2L-2) or control shRNA (Control) were stained with propidium iodide (PI, 3 μg/mL) and analyzed by flow cytometry to evaluate the cell death rate. Data shown as mean ± SD of n=3 technical replicates. **J** Lipid peroxidation was measured by C11-BODIPY (5μM) in 22Rv1 and Du145 cells with control or VSTM2L shRNAs. Data shown as mean ± SD of n=3 technical replicates. **K** The level of glutathione (GSH) in 22Rv1 and Du145 cells with control or VSTM2L shRNAs. Data shown as mean ± SD of n=3 technical replicates. **L** Western blot analysis of GPX4 and VSTM2L protein levels in VSTM2L knockdown Pca cells or control cells. Protein levels were normalized to total GAPDH. *P* value was determined by unpaired two-tailed Student’s *t*-test (**D, E, F**) and one-way ANOVA (**I, J, K**).

Ferroptosis is an iron-dependent, non-apoptotic form of cell death triggered by extensive lipid peroxidation that overwhelms lipid protection mechanisms^30, 31^. To determine whether VSTM2L plays roles in regulating ferroptosis, we performed propidium iodide (PI), Annexin V and BODIPY™ 581/591 C11 (C11-BODIPY) staining experiments to test the effect of VSTM2L suppression on the cell death and lipid peroxidation, respectively. The results indicated that reduced VSTM2L protein levels in 22Rv1 or Du145 cells could increase cell death rate (Fig. 3G-I) and lipid peroxide accumulation (Fig. 3J), but have no obvious effect on phosphatidylserine exposure of Pca cells (Fig. S3 D), suggesting that VSTM2L knockdown in PCa cells induced ferroptosis not apoptosis. To further confirm the regulation of VSTM2L in ferroptosis, we measured intracellular glutathione (GSH) levels in VSTM2L knocked down 22Rv1 or Du145 cells. As expected, down-regulation of VSTM2L in PCa cell lines attenuated GSH levels (Fig. 3K). We also checked the protein levels of the glutathione peroxidase 4 (GPX4), Bcl-2 associated X protein (Bax) and B-cell lymphoma-2 protein (Bcl-2) in VSTM2L knocked down 22Rv1 or Du145 cells by western blot. The results showed that inhibition of VSTM2L expression diminished the protein levels of GPX4 as expected in these two cell lines (Fig. 3L), but have no obvious effect on the level of Bax and Bcl-2 (Fig. S3 E). Furthermore, ferroptosis inhibitor ferrostatin-1(Fer-1) can rescue the cell death caused by knockdown of VSTM2L (Fig. S3 F). Accordingly, overexpression of VSTM2L in PCa cell lines could reverse cell death and the levels of GSH and GPX4 (Fig. S4 A-F). Collectively, these data suggested that VSTM2L acts as a potential suppressor of ferroptosis not apoptosis by GPX4 inhibition in prostate cancer cells.

### VSTM2L inhibition sensitizes PCa cells to ferroptosis inducer, RSL3

To advance our understanding of the role and mechanism of VSTM2L in regulating ferroptosis, we investigated the effect of VSTM2L suppression on ferroptosis sensitivity to RSL3, a small molecule inhibitor target GPX4^32^. Then, cell viability was checked by CCK-8 assay. After 24 hours of RSL3 treatment, the viability of PCa cells with VSTM2L silencing, as indicated by the IC50 values, was significantly reduced compared to that of control cells (Fig. 4A). To further evaluate whether VSTM2L-knockdown promoted RSL3-induced ferroptosis *in vivo*, we constructed a xenograft model with nude mice. A schematic of the *in vivo* experiment, including tumor inoculation, RSL3 injection and the experimental timeline, was shown in Fig. 4B. All experimental mice maintained a good appetite and behavioral status after tumor inoculation and RSL3 administration until they were humanely euthanized and isolated the subcutaneous tumors. As expected, the VSTM2L suppression groups exhibited tumors with reduced volume and weight compared with the control groups, while there was an enhancement in the anti-tumor effects once VSTM2L suppression and RSL3 treatment were combined (Fig. 4C-D). The results suggested that inhibition of VSTM2L expression enhanced the *in vivo* therapeutic efficacy of RSL3. Meanwhile, we also tested the in vivo effects of the combined treatment on GSH level. Consistent with the *in vitro* cell results, VSTM2L knockdown in resected tumors weakened the levels of GSH, which was further reduced by RSL3 administration (Fig. 4E).

**Fig. 4.**
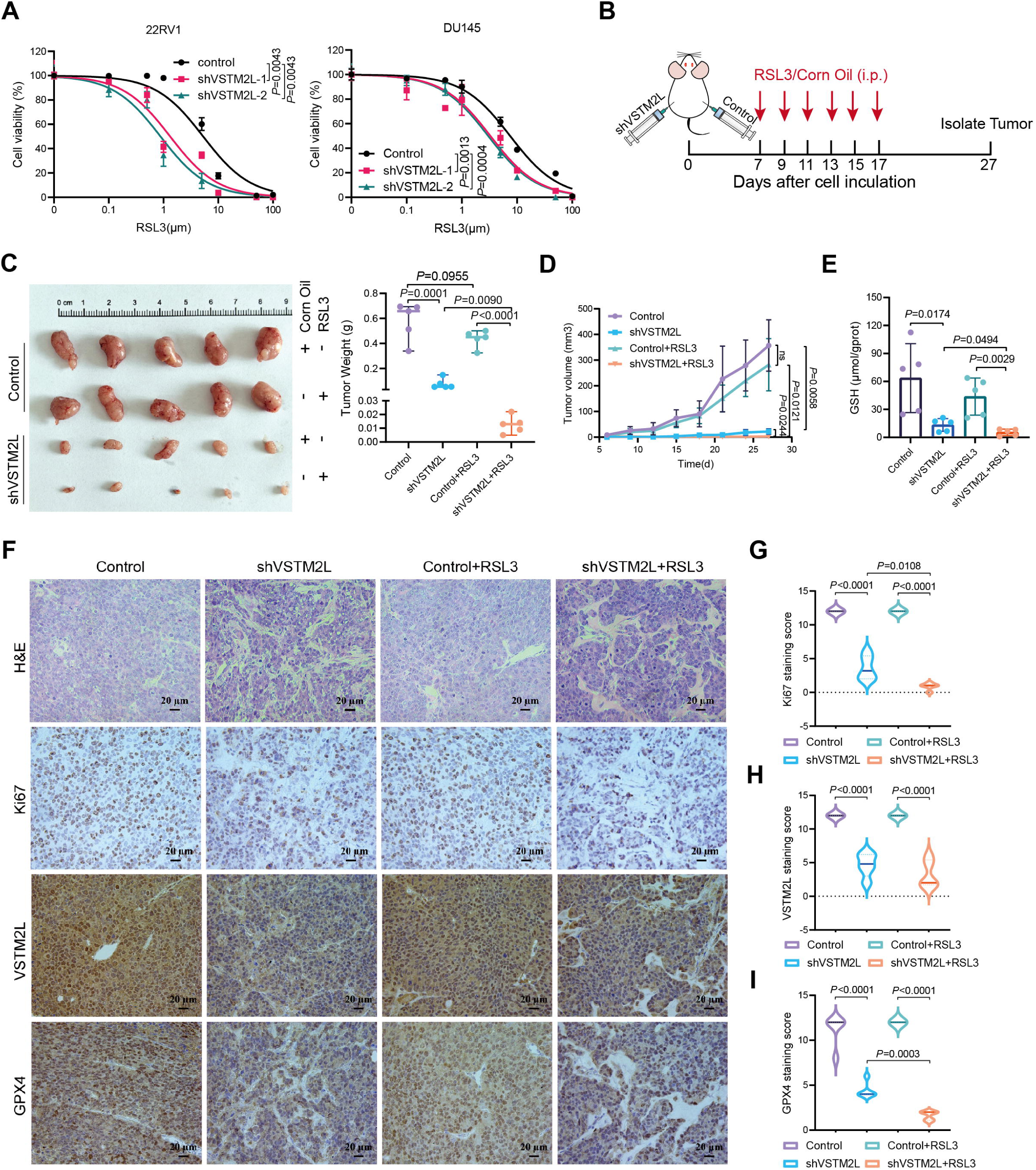
VSTM2L inhibition sensitizes PCa cells to ferroptosis inducer, RSL3. **A** Viability of 22Rv1 or DU145 cells transfected with VSTM2L shRNA or control shRNA was measured by CCK-8 in the presence of varying concentrations of RSL3 (0-100 μM) for 24 h. Data shown as mean ± SD of n=3 technical replicates. **B** The schematic diagram of the *in vivo* experimental process. 1 × 10^6^ 22Rv1 shVSTM2L cells and control cells were inoculated trans-subcutaneous into nude mice. RSL3 (5mg/kg) was intraperitoneally injected into the mice on alternate days for a total of six injections. The tumor tissues were isolated on 27th days post-inoculation. **C, D** General view of tumor weight (**C**) and tumor volume (**D**) (examined every 3 days) of each indicated group at 27 days after cell injection. Data shown as mean ± SD, n=5 tumors, ns, not significant. **E** The levels of GSH in the tumor tissues at the indicated endpoint. Data shown as mean ± SD, n=5 tumors. **F-I** Immunohistochemistry (IHC) and hematoxylin and eosin (H & E) staining for Ki-67 (**F**, **G**), VSTM2 (**F, H**) and GPX4 (**F, I**) were performed in isolated tumor tissues. Data shown as mean ± SD, n=5 randomly selected magnification fields, Scale bar =20 μm. *P* value was determined by two-way ANOVA (**A, D**) and unpaired two-tailed Student’s *t*-test (**C, E, G-I**).

Next, hematoxylin and eosin (H&E) staining was applied to evaluate the histomorphological changes within the tumor tissues. The results revealed that VSTM2L suppression induced cell death, and administration of RSL3 further exacerbated the phenotype (Fig. 4F). Then, immunohistochemical (IHC) staining was utilized to assess the protein levels of VSTM2L, Ki67, and GPX4 in xenograft samples. The results showed that tumors from VSTM2L knockdown group exhibited reduced expression of Ki67, which was lower in the combination group with VSTM2L attenuation and RSL3 treatment, as expected (Fig. 4F, G). Furthermore, the down-regulation of VSTM2L expression was associated with a substantial decrease in GPX4 protein levels in the tumor tissues and the administration of RSL3 further suppressed the expression levels of GPX4 (Fig. 4F, H, I). Thus, these results provided evidence that targeting VSTM2L and inducing ferroptosis may act as an efficient strategy in PCa treatment.

### Fer-1 reverses VSTM2L suppression induced ferroptosis of PCa cells in the presence of RSL3

To further investigate the relationship between VSTM2L and ferroptosis, we performed CCK-8 assay to detect cell viability. The results showed that RSL3-induced cell death in DU145 shVSTM2L cells was abolished by the ferroptosis inhibitor ferrostatin-1(Fer-1), rather than a panel of apoptosis inhibitors (Z-VAD-FMK, Z-IETD-FMK and Z-DEVD-FMK), as well as autophagy inhibitors 3-Methyladenine (3MA) and Chloroquine (CQ) (Fig. 5A). Moreover, VSTM2L knockdown markedly increased RSL3-induced cell death in 22Rv1 cells, and this effect was blocked by Fer-1 (Fig. 5B). We also observed changes in cell morphology after RSL3 and Fer-1 treatments (Fig. 5C). In addition, knockdown of VSTM2L in PCa cells accelerated the increase in RSL3-induced cell death, which was inhibited by Fer-1 (Fig. 5D-F). We also observed that the increased accumulation of lipid peroxidation triggered by RSL3 treatment in VSTM2L suppressed PCa cells could be fully reversed by Fer-1(Fig. 5G, H).

**Fig. 5.**
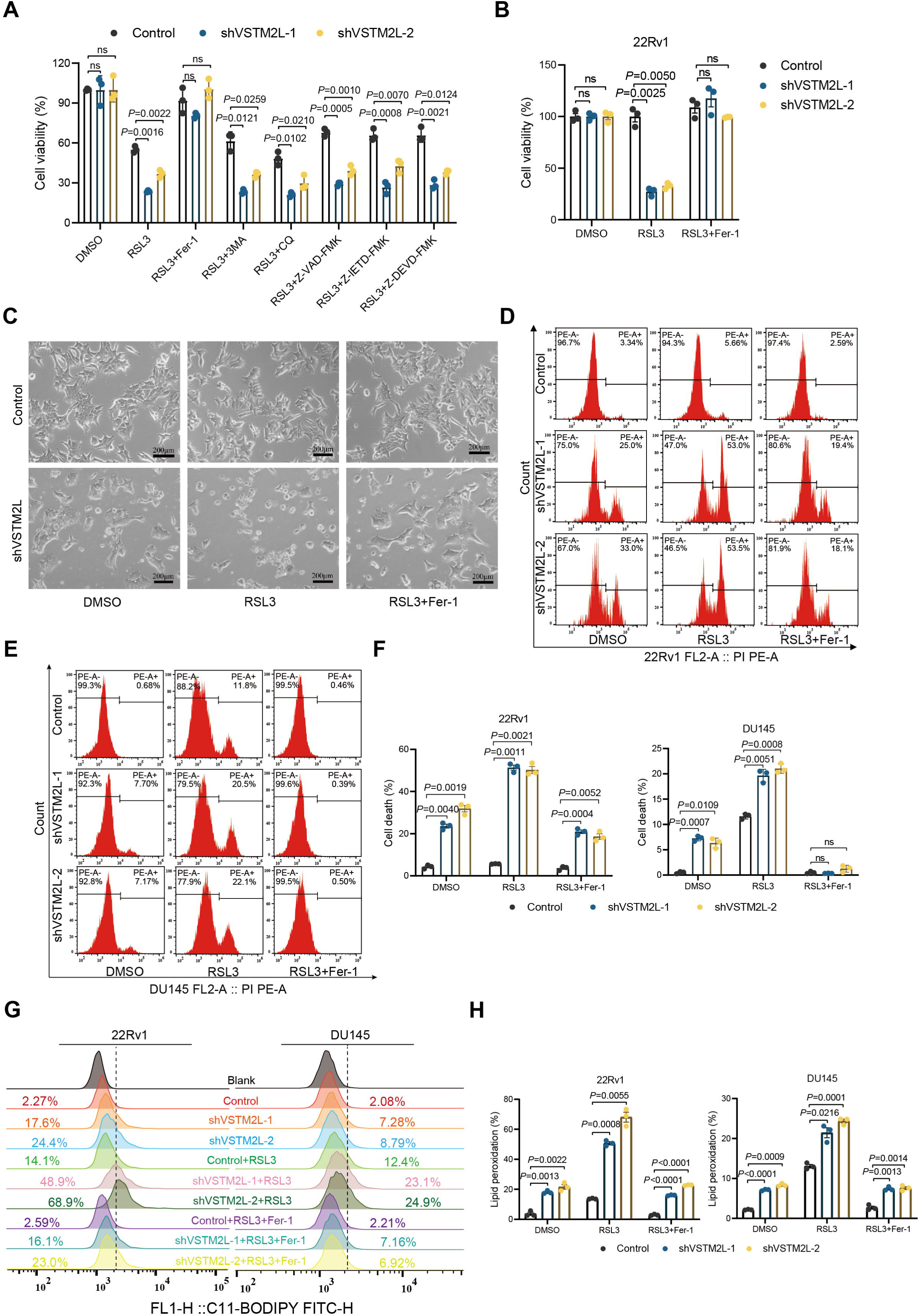
Fer-1 reverses VSTM2L suppression induced ferroptosis of PCa cells in the presence of RSL3. **A** Viability of DU145 cells transfected with VSTM2L shRNA (shVSTM2L-1, shVSTM2L-2) or control shRNA (Control) treated with or without RSL3 (2 μM), Fer-1 (5 μM), 3MA (60 μM), CQ (25 μM), Z-VAD-FMK (10 μM), Z-IETD-FMK (10 μM), Z-DEVD-FMK (10 μM) for 24h. Data shown as mean ± SD, n=3, ns, not significant. **B** Viability of 22Rv1 cells transfected with VSTM2L shRNAs or control shRNA treated with or without RSL3 (1 μM), Fer-1(10 μM). Data shown as mean ± SD, n=3, ns, not significant. **C** Bright-field images of 22Rv1 cells with VSTM2L or control shRNA treated with or without RSL3 (1 μM) or Fer-1 (10 μM) for 24h. **D-F** Representative flow cytometry histograms show the percentage of cell death in 22Rv1 (**D**) and DU145 (**E**) cells transfected with VSTM2L shRNAs or control shRNA, stained by PI (3 μg/mL). Quantification of the percentage of cell death rate in different cell lines (**F**). Cells were treated with or without RSL3 (DU145, 2 μM; 22Rv1, 1 μM) and Fer-1 (DU145, 5 μM; 22Rv1, 10μM) for 24h. Data shown as mean ± SD, n=3, ns not significant. **G** Relative C11-BODIPY fluorescence measured by flow cytometry of PCa cells treated with VSTM2L shRNA or control shRNA and cultured with either RSL3 (DU145, 2 μM; 22Rv1, 1 μM), Fer-1 (DU145, 5 μM; 22Rv1, 10μM) or both for 24 h. **H** The percentages of lipid peroxidation are presented as the means ± SD, n = 3. ns, not significant. *P* value was determined by two-way ANOVA.

In summary, VSTM2L ablation promotes RSL3 induced ferroptosis of PCa cells, which was blocked by Fer-1. These findings provided robust support for VSTM2L as a potential target to ferroptosis in prostate cancer.

### VSTM2L suppresses VDAC1 oligomerization by facilitating the interaction between VDAC1 and HK2

VDAC1, located in the outer membrane of mitochondria (OMM), serves as a crucial gene for mitochondria quality control^33^. The oligomers of VDAC1 plays an important role in maintaining mitochondrial function and overall cell viability, including import phospholipids into mitochondria^34^. A recent published study indicates that phospholipids promote ferroptosis in mammalian cells^7^. To further investigate the mechanism of VSTM2L in regulating ferroptosis via interacting with VDAC1, we co-transfected distinct tagged versions of VDAC1 (Myc- and Flag-tagged) and different amounts of 3 × Flag-VSTM2L vectors into HEK293T cells. Subsequently, we conducted co-IP assay using anti-Myc antibody. The results suggested that the increased amount of VSTM2L did not affect the protein levels of VDAC1, but disturbed the oligomerization between Myc-VDAC1 and Flag-VDAC1, indicating that forced expression of VSTM2L could inhibit VDAC1 oligomerization (Fig. 6A). To further confirm this finding, we performed a cross-linking assay using ethylene glycol bis-(succinimidyl succinate) (EGS). Notably, the suppression of VSTM2L promotes the formation of VDAC1 oligomers; accordingly, VSTM2L overexpression inhibits the oligomerization of VDAC1 in prostate cancer cells (Fig. 6B).

**Fig. 6.**
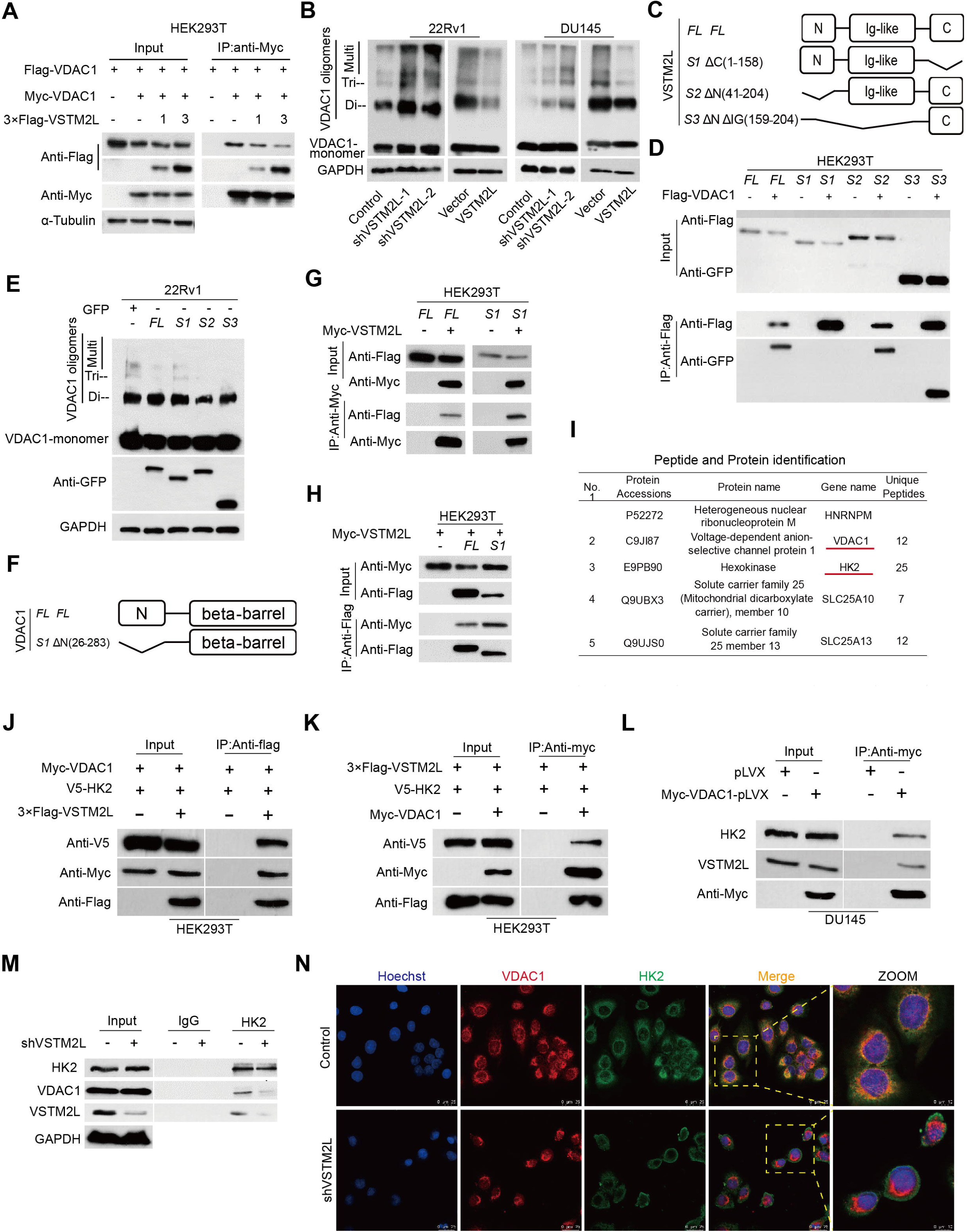
VSTM2L suppresses VDAC1 oligomerization by facilitating the interaction between VDAC1 and HK2. **A** Co-transfected Myc-VDAC1, Flag-VDAC1 and different amounts of 3×Flag-VSTM2L into HEK293T cells, the oligomerization of VDAC1 was analyzed by co-IP/WB. **B** VSTM2L suppressed or overexpressed PCa cells were harvested, respectively, and incubated with ethylene glycol bis-(succinimidyl succinate) to cross-link proteins, then subject to western blot to evaluate VDAC1 oligomeric status. **C** Schematic representation of the full-length VSTM2L and its truncated mutants was shown. **D** HEK293T cells were transfected with indicated full-length VSTM2L or its truncated forms, as well as full-length VDAC1. Cell lysates were harvested and subjected to immunoprecipitation with anti-Flag antibodies to map the binding region of VDAC1 with VSTM2L. **E** 22Rv1 cells were transfected with full-length VSTM2L or its truncated mutant as indicated. Cell lysates were collected and analyzed using a cross-linking assay to identify the functional region of VSTM2L responsible for inhibiting VDAC1 oligomerization. **F** A schematic of the full-length VDAC1 and its N-terminus truncated mutant was presented. **G, H** HEK293T cells were co-transfected with indicated full-length or N-terminus truncated mutant of VDAC1 and full-length VSTM2L. Cell lysates were collected and subjected to immunoprecipitation with anti-Myc (**G**) or anti-Flag (**H**) antibodies. Immunoprecipitants were analyzed with anti-Flag antibody and anti-Myc antibody. **I** A partial list of binding proteins of VSTM2L in LNCaP cells identified by mass spectrometry analysis. **J, K** HEK293T cells were co-transfected with V5-HK2, Myc-VDAC1 and 3×Flag VSTM2L. Cell lysates were harvested and subjected to immunoprecipitation with anti-Flag (J) and anti-Myc (K) antibodies. Immunoprecipitants were analyzed with anti-Flag antibody, anti-V5 antibody and anti-Myc antibody. **L** DU145 cells overexpressed Myc-VDAC1-pLVX or control cells were subjected to immunoprecipitation with anti-Myc antibody and followed by western blot analysis. **M** The whole cell lysates were prepared from DU145 cells transfected with VSTM2L shRNA lentivirus or control shRNA lentivirus, and immunoprecipitations were performed with anti-HK2 antibodies, followed by western blot with indicated antibodies. **N** Representative images of immunofluorescence staining for VDAC1 and HK2 in DU145 cells. The nucleus was stained with Hoechst. Scale bar 25 μm and 10 μm.

To explore the mechanism of VSTM2L in inhibiting the oligomerization of VDAC1, we established a series of functional domain deletions of VSTM2L (Fig. 6C). Mapping the binding domains revealed that the C-terminal domain of VSTM2L was essential for the binding to VDAC1 (Fig. 6D). Afterwards, we transfected the full-length (*FL*) and variant of functional domain deletions of VSTM2L into 22Rv1 cells. Cross-linking assay confirmed that full-length VSTM2L and its two domain deletions (*S2*, *S3*) indeed apparently inhibited the oligomerization of VDAC1, instead of C-terminal deletion (*S1*) of VSTM2L (Fig. 6E).

It was reported that the N-terminus of VDAC1 is the functional domain of VDAC1^35^. And the N-terminus of VDAC1 is crucial for VDAC1 oligomerization^36, 37^. Thus, the truncated mutant of N-terminus of VDAC1 (*S1*) was constructed to check whether VSTM2L could bind to N-terminus of VDAC1 by immunoprecipitation (Fig. 6F). The results showed that VSTM2L can bind at the C-terminus of VDAC1 (Fig. 6G-H). These findings prompted us to hypothesize that the inhibition of VDAC1 oligomer formation by VSTM2L is involved in participation of other proteins. To confirm this hypothesis, we checked the proteins interacted with VSTM2L by IP/MS analysis, and found that Hexokinase 2 (HK2) might be the potential protein connecting these two factors(Fig. 6I), as HK2 has been well-documented for the interaction with the N-terminus of VDAC1 to constrain VDAC1 oligomerization^36, 38, 39^. Importantly, our exogenous and semi-endogenous co-IP assay suggested that a tripartite complex comprising VSTM2L, VDAC1, and HK2 indeed exists (Fig. 6J-L). This observation raised the question that whether VSTM2L inhibited VDAC1 oligomerization through facilitating the binding affinity between VDAC1 and HK2. To address this question, we utilized DU145 cell lysates with VSTM2L knockdown and performed immunoprecipitation with HK2-specific antibodies in parallel with IgG control antibodies to capture VDAC1 protein. The results showed that knockdown of VSTM2L weakened the interaction between VDAC1 and HK2 (Fig. 6M). What’s more, immunofluorescence staining showed that compared with the control group, HK2 was dissociated from VDAC1 after knockdown of VSTM2L in prostate cancer cells (Fig. 6N). Collectively, the results mentioned above indicated that VSTM2L was essential for mediating the interaction between VDAC1 and HK2.

In conclusion, these findings substantiated our hypothesis that VSTM2L enhances the binding affinity between VDAC1 and HK2, thereby inhibiting the oligomerization of VDAC1.

### VSTM2L suppresses ferroptosis and maintains mitochondria homeostasis via inhibiting VDAC1 oligomerization in PCa cells

Accumulated evidences indicated that mitochondria homeostasis is crucial for suppression of ferroptosis in cancer cells^7, 10, 40, 41^, VDAC1 oligomerization play a pivotal role in regulation of ferroptosis via maintaining the balance of mitochondrial reactive oxygen species (mtROS)^26^. Mitochondiral ROS was reported to induce ferroptosis^40, 42, 43^, while the mechanism by which ferroptosis is triggered in mitochondria is still unknown. Our quantification of mtROS test using MitoSOX probe, revealed that suppression of VSTM2L markedly increased mtROS levels in PCa cells compared with control group (Fig. 7A-B). Accordingly, overexpression of VSTM2L in PCa cells led to the decreased mtROS levels (Fig. S5 A).

**Fig. 7.**
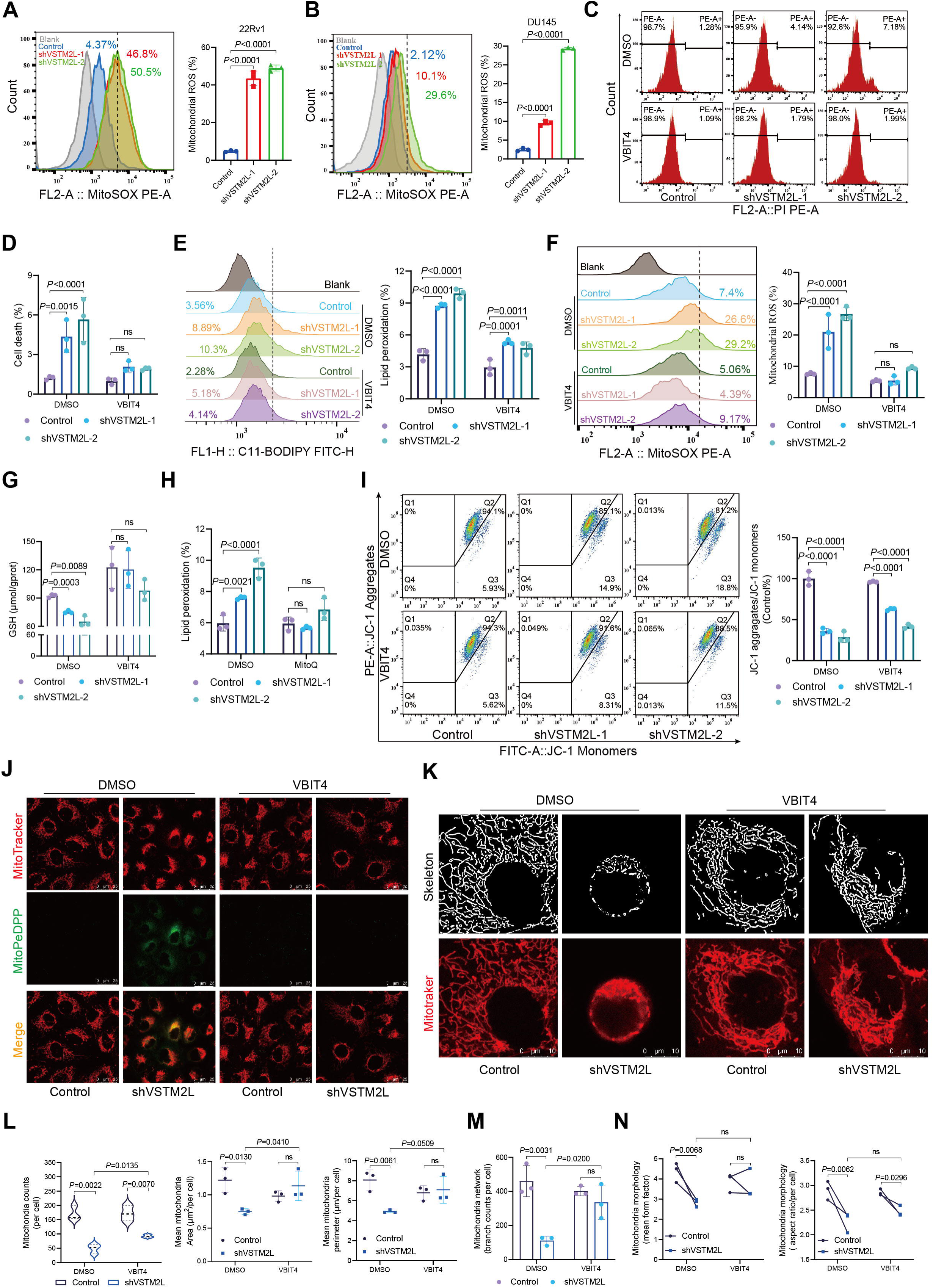
VSTM2L suppresses ferroptosis and maintains mitochondria homeostasis via inhibiting VDAC1 oligomerization in PCa cells. **A, B** The levels of mtROS in 22Rv1 (**A**) and DU145cells (**B**) transfected with VSTM2L shRNA (shVSTM2L-1, shVSTM2L-2) or control shRNA (Control). **C-F** VSTM2L knockdown DU145 cells were cultured with or without VBIT4 (5 μM) for 24 h, then, cell death (**C, D**), lipid peroxidation (**E**) and mtROS (**F**) were assessed by PI (3 μg/μL), C11-BODIPY (5 μM) and MitoSOX (5 μM) staining, respectively. **G** The level of GSH in VSTM2L inhibited DU145 cells treated with or without VBIT4 (5 μM) for 24h. **H** The level of lipid peroxidation in VSTM2L suppressed DU145 cells cultured with or without MitoQ (10 nM) for 24h and assessed by C11-BODIPY (5 μM) staining. **I** The level of MMP in VSTM2L knockdown DU145 cells cultured with or without VBIT4 (5 μM) for 24h. **J, K** Representative images of mitochondrial in VSTM2L inhibited DU145 cells cultured with or without VBIT4 (5 μM) by MitoPeDPP (0.5 μM, Scale bars = 25 μm) (**J**) and Mito-Tracker (200 nM, Scale bars = 10 μm) (**K**) staining. **L-N** Mitochondria counts, mean mitochondria area, mean mitochondria perimeter (**L**), mitochondria network (branch counts) (**M**), mitochondria morphology (mean mitochondria form factor and mean mitochondria aspect ratio) (**N**) for images shown in **K** by ImageJ were determined. Data shown as mean ± SD, n=3 biologically independent experiments. *P* value was determined by one-way (**A**) and two-way ANOVA (**D-I, L-N**).

To explore whether VSTM2L suppression contributed to the ferroptosis of PCa cells through regulating oligomerization of VDAC1, we employed a VDAC1 oligomerization inhibitor, VBIT4^44, 45^, in our subsequent studies. The increased cell death, massive lipid peroxidation accumulation, elevated mtROS levels and diminished GSH concentrations, caused by VSTM2L knockdown were dramatically reversed by VBIT-4 (Fig. 7C-G; Fig. S5 B-C). Further, the shVSTM2L cells were cultured with a mitochondira-targeted antioxidant, Mitoquinone (MitoQ), which significantly reduced the accumulation of lipid peroxidation and mtROS (Fig. 7H; Fig. S5 D). These findings suggested that VDAC1 oligomerization inhibitor prevented PCa cells from VSTM2L knockdown-induced VDAC1 oligomerization and ferroptosis. And the accumulation of mtROS, resultant from increased formation of VDAC1 oligomers, is the principal cause of ferroptosis in VSTM2L inhibition PCa cells.

To further confirm the notion that the promotion of VDAC1 oligomerization, consequent to VSTM2L knockdown, triggers a comprehensive collapse of mitochondrial homeostasis in PCa cell lines, the JC-1 probe was applied to assess MMP. The results illustrated an elevation in MMP levels following the suppression of VSTM2L, and this effect was counteracted by VBIT4 (Fig. 7I). It has been established that mitochondrial lipid peroxidation, driven by accumulated mtROS, plays a pivotal role in the initiation of ferroptosis^46^. Herein, we employed MitoPeDPP^47^ probe to check mitochondrial lipid peroxidation detected by using a confocal microscopy. The results revealed that VSTM2L knockdown in DU145 cells showed pronounced bright green fluorescence compared with the control cells, and the MitoPeDPP signal caused by VSTM2L suppression was virtually abolished in the presence of VBIT4 (Fig. 7J). Notably, living imaging of subcellular mitochondria by confocal microscopy demonstrated that VSTM2L knockdown in DU145 cells led to a dramatic mitochondrial morphology and network changes, including reductions in mitochondria number, mean mitochondria area, mean mitochondria perimeter, and branch counts, which were reversed by VBIT4 (Fig. 7K-M; Fig. S6). However, VBIT4 did not show any significant improvement on the reduction of mean mitochondria form factor and mean mitochondria aspect ratio, which were induced by the downregulation of VSTM2L in DU145 cells (Fig. 7N). The results explained the reason that VBIT4 cannot fully reverse the various changes brought about by the knockdown of VSTM2L in PCa cells.

Collectively, the aforementioned results provided ample evidences that the enhanced VDAC1 oligomerization caused by VSTM2L knockdown led to a disruption of mitochondrial homeostasis, thereby contributing to the ferroptosis of PCa cells.

### VSTM2L plays oncogenic roles in PCa by inhibiting VDAC1 oligomerization

To explore whether the oligomerization of VDAC1 suppressed by VSTM2L contribute to regulating PCa cells growth, we conducted experiments both *in vitro* and *in vivo* with the intervention of VBIT4. We first cultured the shVSTM2L DU145 cells or control cells under treatment with VBIT4 for 24 hours. EGS-based cross-linking assay suggested that VBIT4 successfully reversed the oligomerization of VDAC1 promoted by the suppression of VSTM2L (Fig. 8A). Then, CCK-8 assay declared that VBIT4 could obviously rescue the diminished proliferation and colony formation abilities caused by VSTM2L knockdown in DU145 cells (Fig. 8B-D). Additionally, VBIT4 reversed the reduced migration rate in shVSTM2L cells as well (Fig. 8E-H). Moreover, 3.5 × 10^6^ DU145 shVSTM2L cells and the control cells were inoculated trans-subcutaneous into nude mice to establish a xenograft model, respectively, the schematic of the experiment was shown in Fig. 8I. VBIT4 (20mg/kg) was exposed to the mice by gavage each other day until the endpoint. Importantly, VBIT4 had demonstrated the capacity to ameliorate the decelerated growth of tumor tissues resulting from VSTM2L knockdown within the tumor tissues (Fig. 8J-L). The IHC and H&E staining results further indicated that the administration of VBIT4 to experimental animals notably reversed the ferroptotic cells death within the tumor tissues caused by VSTM2L suppression (Fig. 8M-N). Meanwhile, VSTM2L knockdown in tumor tissues reduced the expression of Ki-67, and this effect was blocked by VBIT4 (Fig. 8M, O).

**Fig. 8.**
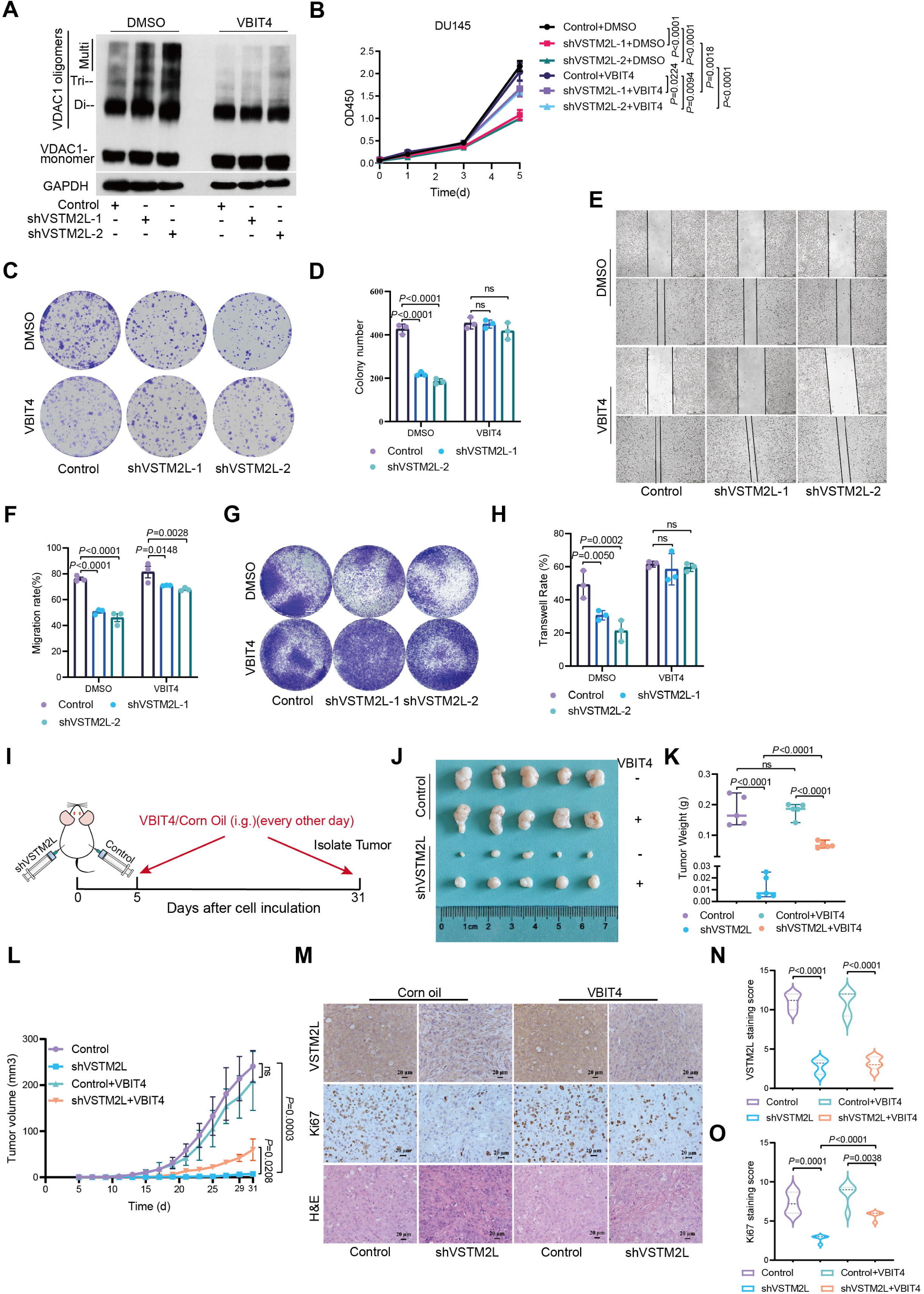
VSTM2L plays oncogenic roles in PCa by inhibiting VDAC1 oligomerization. **A** VSTM2L knockdown DU145 cells cultured with or without VBIT4 (5μM) for 24 h were harvested for the subsequent cross-link assay, then subject to western blot to assess VDAC1 oligomeric status. **B-D** VSTM2L suppressed DU145 cells were cultured with or without VBIT4 (5μM) for the indicated time, then the cell viability (**B**) and colony formation ability (**C, D**) were analyzed by CCK8 assay and Crystal Violet Aqueous Solution staining, respectively. Data shown as mean ± SD, n=3, ns not significant. **E, F** Wound healing analysis of VSTM2L inhibited DU145 cells in the absence or presence of VBIT4 (5μM). Representative images (**E**) and quantification (**F**) are shown as indicated. **G, H** Representative pictures (**G**) and quantification analysis (**H**) of migration assays in VSTM2L knockdown DU145 cells cultured with or without VBIT4 (5μM). Data shown as mean ± SD, n=3, ns not significant, scale bars = 250 μm. **I** A schematic diagram of the *in vivo* experimental process. 3.5 × 10^6^ DU145 shVSTM2L cells and control cells were inoculated trans-subcutaneous into nude mice. The mice were administrated with VBIT4 (20mg/kg) by gavage on alternate days until the tumor tissues were isolated on 31th days post-inoculation. **J, K** General view of tumor weight of each indicated groups at the endpoint. Data shown as mean ± SD, n=5 tumors, ns not significant. **L** Tumor growth curves were shown. Data shown as mean ± SD, n=5 tumors, ns not significant. **M-O** Immunohistochemistry (IHC) and hematoxylin and eosin (H & E) staining for VSTM2L (**M, N**) and Ki-67 (**M, O**) were performed in isolated tumor tissues. Data shown as mean ± SD, n=5 randomly selected magnification fields, Scale bar =20 μm. *P* value was determined by unpaired two-tailed Student’s *t*-test (**K, N, O**) and two-way ANOVA (**B, D, F, H, L**).

Taken together, these results indicated that VSTM2L exerts its oncogenic roles in prostate cancer by impeding the assembly of VDAC1 oligomers both *in vitro* and *in vivo*.

## DISCUSSION

In the present study, we found that VSTM2L, a VDAC1 binding partner, located in mitochondria plays an oncogenic role in PCa cells. Suppression of VSTM2L led to the dissociation of HK2 from VDAC1 and the increased oligomerization of VDAC1, which disturbed the homeostasis of mitochondria, triggered the ferroptosis and eventually restrained PCa progression (Fig. 9).

**Fig. 9.**
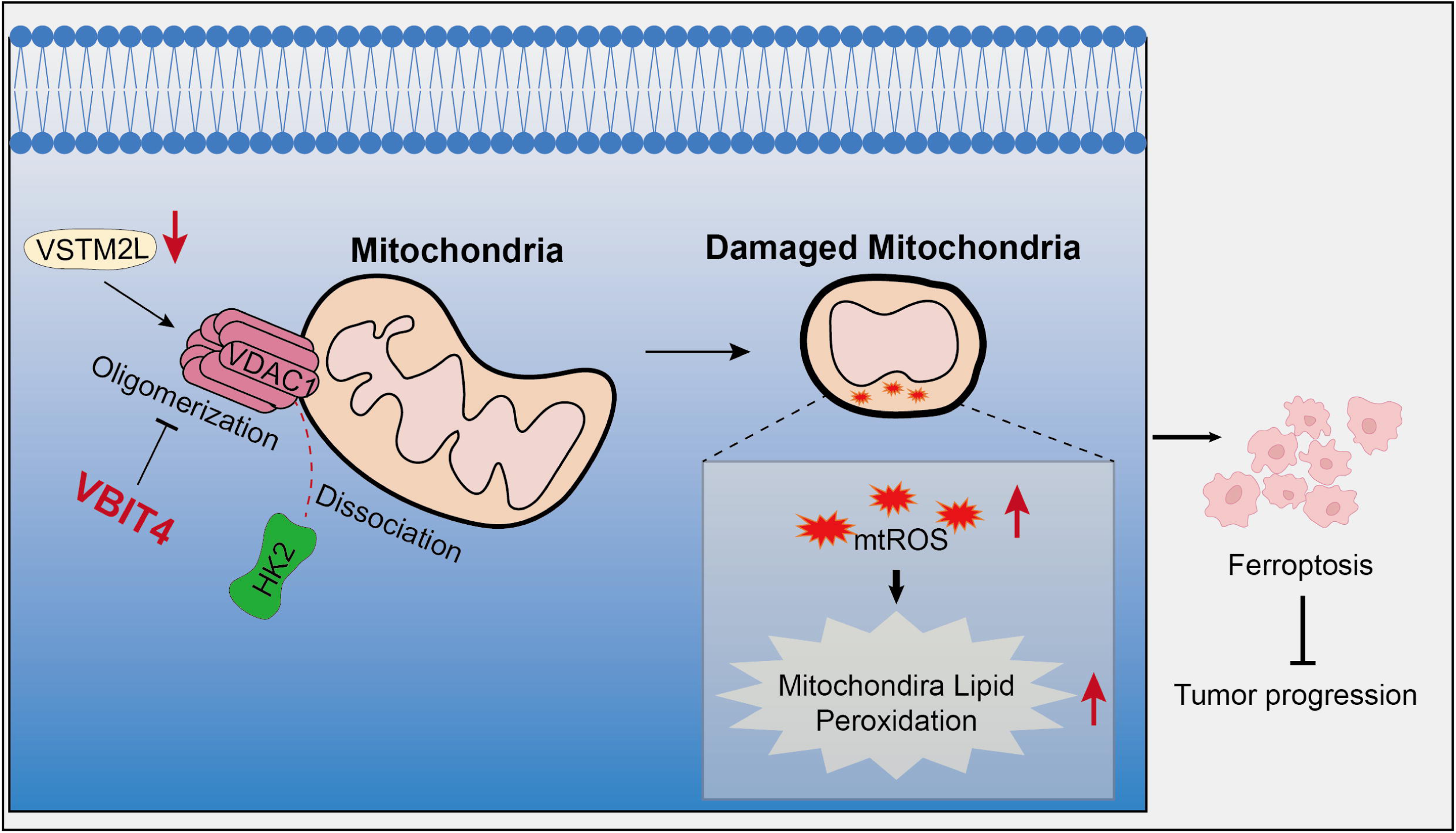
Schematic diagram showing that VSTM2L knockdown facilitates VDAC1 oligomerization, disturbs the homeostasis of mitochondria and triggers ferroptosis. Suppression of VSTM2L led to the dissociation of HK2 from VDAC1 and the increased oligomerization of VDAC1, which disturbed the homeostasis of mitochondria, ultimately triggering ferroptosis and inhibiting PCa progression.

In recent years, the roles of VSTM2L in different cancers have already been reported in the literature. It was reported that VSTM2L is recognized as a potential biomarker associated with ovarian cancer metastasis and prognosis^28^. High expression of VSTM2L induced resistance to chemo radiotherapy in rectal cancer and might be a potential resistant predictable biomarker for advanced rectal cancer patients receiving preoperative chemo radiotherapy^29^. Recently, VSTM2L was found related to the viability and survival of cholangiocarcinoma cells^48^. So far, the function of VSTM2L in PCa progression and its location in Pca cells has never been reported. Our study suggested that VSTM2L was highly expressed in prostate tumors and positively associated with the poor prognosis of PCa patients. Meanwhile, we reported that VSTM2L is a mainly mitochondria located protein in Pca cells. Suppression of VSTM2L weakened cell growth and migration ability in PCa. Our findings indicated that VSTM2L plays an oncogenic role in PCa progression.

Ferroptosis is a novel form of programmed cell death dependent on iron and reactive oxygen species, is mainly characterized by mitochondrial shrinkage, increased density of mitochondrial membranes, reduction or vanishing of mitochondria crista, the accumulation of lipid peroxidation and GSH depletion^49, 50, 51^. In our study, we observed a remarkable shrinkage of the cells with shRNA targeting VSTM2L. We also found significant alterations of the morphology of mitochondria in VSTM2L knocked down 22Rv1 cells, mainly characterized by a general shrinkage in size, increased membrane density, and diminution in cristae. Reduction of VSTM2L levels in PCa cells led to increased cell death, lipid peroxide accumulation and decreased GSH levels. In addition, VSTM2L knockdown in PCa cells restrained the protein levels of GPX4. These findings suggested that VSTM2L is a novel modulator of ferroptosis and protects PCa cells against ferroptosis. However, how VSTM2L regulates the GPX4 expression need to be further investigation.

Accumulating evidences indicated that RSL3 can induce ferroptosis by inhibiting GPX4 and suppress tumor growth^52, 53^. Ferroptosis inducers (Erastin or RSL3) and in combination with standard-of-care second-generation antiandrogens are novel therapeutic strategies for advanced prostate cancer^54^. Ferroptosis has been confirmed to enhance the sensitivity of castration-resistant prostate cancer cells to therapeutic agents^5, 6^. In this study, we discovered that VSTM2L knockdown enhanced the sensibility of RSL3-induced ferroptosis in vitro and in vivo, the discovery holds significant implications for VSTM2L as a novel prognostic target for prostate cancer.

VDAC1 plays a crucial role in regulation of ferroptosis, and this regulation is achieved through the oligomeric form of VDAC1 rather than its monomeric form. It was reported that protecting mitochondria via inhibiting VDAC1 oligomerization weakened hepatocyte ferroptosis^26^. Zhou *et al*. reported that BABP31 could directly bind to VDAC1 and regulate gastric cancer cells ferroptosis through affecting VDAC1 stability and oligomerization^25^. However, the mechanisms of VDAC1 oligomerization in regulating ferroptosis are not well characterized. Our study found that VSTM2L knockdown in PCa cells promoted VDAC1 oligomerization and inhibited cell growth and migration, which were reversed by VBIT4, an inhibitor of VDAC1 oligomerization, *in vitro* and *in vivo*. These findings indicated that VSTM2L regulated prostate cancer progression and ferroptosis via directly binding to VDAC1 and affecting VDAC1 oligomerization.

The N-terminus of VDAC1 has been identified as the functional domain and response for the oligomerization of VDAC1^35, 36, 37^. Our results suggested that VSTM2L mainly bind to the C-terminus other than N-terminus of VDAC1. Interestingly, we discovered a novel binding protein of VSTM2L, HK2, which is well-documented in the literature for its interaction with the N-terminus of VDAC1^36, 38, 39^, thereby constraining VDAC1 oligomerization^24^. Shangguan *et al*. implicated HK2 in the promotion of prostate cancer progression, primarily through its interaction with the N-terminal of VDAC1, which limits oligomers formation^55^. Here, we discovered that VSTM2L can form a complex with HK2 and VDAC1, and facilitate the binding of HK2 to VDAC1, thereby inhibiting ferroptosis of PCa.

Mitochondrial homeostasis is increasingly acknowledged as essential players in initiating and amplifying ferroptosis^7, 8^. Shoshan-Barmatz *et al*. reported that VDAC1 acted as a key role in regulating mitochondrial homeostasis and cell death^13, 38, 45^, mainly relies on the interactions with the relative proteins. Yagoda et al. reported that erastin, the first ferroptosis activator identified in 2003, binds to VDAC2/3 to change mitochondrial membrane permeability and reduce NADH oxidation rates, a mechanism distinctly different from how VDAC1 regulates ferroptosis^56^. Herein, we found that VSTM2L suppression led to the increased cell death, diminished GSH levels, the increased MMP levels, mitochondrial lipid peroxidation accumulation, elevated mtROS levels and changed morphology of mitochondria, which were all blocked by VBIT4. Importantly, MitoQ reduced the accumulation of lipid peroxidation and mtROS caused by VSTM2L knockdown. These results suggested that the enhanced VDAC1 oligomerization by VSTM2L knockdown led to a disruption of mitochondrial homeostasis. And mtROS burst induced by the upregulated VDAC1 oligomerization, contributed to mitochondrial homeostasis disruption and then onset of PCa ferroptosis. These findings were consistent with the previous study that mtROS imbalance is accountable for disrupting mitochondrial homeostasis and triggering ferroptosis^40, 42, 43^. In summary, VDAC1 binding partner VSTM2L was identified as a novel oncogene located in mitochondria and a ferroptosis suppressor in prostate cancer, and VSTM2L regulated PCa progression and ferroptosis via affecting oligomerization of VDAC1. Thus, VSTM2L could serve as a prognostic biomarker for PCa progression and a therapeutic target for PCa treatment.

## MATERIALS AND METHODS

### Cell culture

Human normal prostate epithelial cell line RWPE1, prostate cancer cell lines (DU145, PC3, 22Rv1 and LNCaP) and HEK293T cells were purchased from American Type Culture Collection (ATCC). All cell lines were confirmed to be mycoplasma free during our study. All the cells were cultured at 37 ℃, with 95 % air and 5 % CO_2_. RWPE-1 cells were cultured with keratinocyte serum-free medium (Gibco, # 10724-011) with bovine pituitary extract and human recombinant epidermal growth factor (EGF); HEK293T and DU145 cell lines were cultured in Dulbecco’s modified Eagle’s medium (DMEM, Gibco, # 12800017), PC3 cells were cultured in Ham’s F-12 medium (HyClone, # SH30526.01), 22Rv1 and LNCaP cell lines were cultured in Roswell Park Memorial Institute (RPMI) 1640 medium (Gibco, # 31800022). 10% (Volume/Volume, v/v) fetal bovine serum (FBS, MCE, # HY-T1000) and 1% (Volume/Volume, v/v) penicillin-streptomycin (PS, Solarbio, # P1400) were supplied to the base medium.

### Plasmid constructs and transfection

*VSTM2L*, *VDAC1*, *HK2* cDNAs were generated in our lab and subcloned into pFLAG-CMV-2, pcDNA3.1 Myc-His A, p3×FLAG-CMV-8, pcDNA3.1 V5-His A, pEGFP-N1, pLVX-mCNV-Zsgreen1-puro, as indicated in the corresponding figures. The cDNA of *VSTM2L* was subcloned into the pCDH-CMV-MCS-EF1-Puro vector. Using full-length pEGFP-N1-VSTM2L and p3×FLAG-CMV-8-VDAC1 cDNAs as templates, a series of functional domain deletions of VSTM2L and VDAC1 were generated.

For lentiviral construct, the shRNA constructs targeting *VSTM2L* were designed and inserted into the lentiviral vector pLKO.1, the Myc-VDAC1 cDNA was subcloned to lentiviral vector pLVX-mCNV-Zsgreen1-puro. For lentiviral production, HEK293T cells were co-transfected with indicated shRNA constructs or overexpression constructs (1.5 µg each), pMD2G (envelope plasmid, 0.375 µg) and psPAX2 (packaging plasmid, 1.125 µg). In all transfection experiments, Lipofectamine 3000 reagents (Invitrogen, # L3000015) or PEI (MCE, # HY-K2014) were utilized according to the manufacturer’s instructions. The primer and shRNA sequences mentioned previously were comprehensively enumerated in Supplementary Table 1.

### Xenograft models

All animal experiments in this project were approved by the Animal Care and Use Committee of Shaanxi Normal University. 4 weeks old male BALB/C-nu/nu mice were obtained from HFK Bioscience CO., LTD (Beijing, PR China) and housed in pathogen-free facilities. All mice were randomized, with each group consisting of five mice, subsequently, one week of adaptive feeding was implemented. PCa cell lines were suspended and counted in cold phosphate-buffered saline (PBS), and 1×10^6^ 22Rv1 cells (150μL, contain 30% matrigel) or 3.5×10^6^ DU145 cells (150μL) were injected into mice subcutaneously. When the tumors reached a minimum size of 1×1mm, the mice were assigned randomly into different treatment groups. RSL3 and VBIT4 were dissolved in DMSO and diluted in corn oil. RSL3 (5mg/kg) was intraperitoneally injected into the mice each other day, within 10 days. VBIT4 (20mg/kg) was exposed to the mice by gavage each other day until the endpoint. Tumor size was measured by vernier caliper every 3 days and the volume was calculated according to the following formula: volume=length×width^2^×1/2. Mice were sacrificed at the endpoint as indicated in the corresponding figures, meanwhile, the tumors were isolated and saved in 4% paraformaldehyde until use.

### Western blot

Total protein of PCa cell lines was extracted with lysis buffer (500mM NaCl, 20mM Tris-HCL pH 8.0, 1% NP-40, 5mM EDTA, 1mM DTT, 1 × Protease Inhibitor), the protein concentration was measured using Bicinchoninic Acid Protein Assay Kit (Thermo Scientific, # 23227). Then, 80 μg protein samples mixed with loading buffer were loaded and separated by 15% SDS-PAGE gel (Epizyme Biotech, # PG214). Finally, the protein bands were exposed on a film using the ECL kit in a darkened environment. All primary anti-bodies and concentrations used in this study were listed in the Supplementary table 2.

### GST Pull-down assay

GST Pull-down assay was conducted as previously described by our team^57^. Briefly, *VSTM2L* cDNA was cloned to pGEX-6P-1 (GST fusion tag), and the recombinant GST-VSTM2L was expressed in E.coli BL 21, induced by isopropyl-β-D-thiogalactopyranoside (IPTG, 0.5mM final concentration) at 30 ℃overnight on a constant temperature shaker. Subsequently, the bacteria solution was centrifugated at 4000g for 3 min to collect the pellets at the bottom of the tube. After washing the pellets twice with cold PBS, the pellets were then resuspended and lysed in lysis buffer described above. The expression of recombinant GST-VSTM2Lprotein was confirmed by SDS-PAGE, followed by Coomassie Brilliant Blue staining. Furthermore, glutathione-Sepharose 4B beads (Thermo Scientific, # 16101) were washed three times using 1 mL cold PBS, before incubated with GST or GST-fusion protein at 4 ℃. The combined beads were collected and washed as above and then incubated with PC3 cell lysis at 4 ℃ overnight on a rotor. Finally, beads were harvested and washed as above, resuspended in 2 × Loading buffer and probed by western blotting.

### Co-Immunoprecipitation coupled with mass spectrometry (CO-IP/MS)

Cell pellets were harvested and resuspended in 600μL lysis buffer by sonication for 1min, and then centrifuged at 12000g for 15min at 4 ℃. For coimmunoprecipitation, the supernatant was incubated with 3μg indicated antibodies for 8-10 hours at 4 ℃ on a rotor. Subsequently, the combined protein lysis was incubated with 45μL of protein A/G-agarose beads (Thermo Scientific, # 20422) at 4 ℃ on a rotor overnight. The combined beads were washed three times with cold lysis buffer, and mixed with 2 × loading buffer for western blot analysis.

For immunoprecipitation / mass spectrometry (IP/MS), LNCaP cells overexpressing Flag-VDAC1 or 3×Flag-VSTM2L were harvested, resuspended in 700μL of lysis buffer, and centrifuged at 12000g for 15min at 4 ℃. The supernatant was incubated with 7μg anti-Flag antibody for 8-10 hours at 4 ℃, followed by incubating with 70μL of protein A/G-agarose beads at 4 ℃ overnight. After extensive washes, the immunoprecipitates were resolved with 2 × loading buffer for 15 % SDS-PAGE gel separation. After the gel was stained with Coomassie Brilliant Blue, the differential bands were excised for proteomic analysis. The specific conditions for Coomassie Brilliant Blue staining and LC-MS are displayed in Supplementary figure 1, Supplementary File 1, and Supplementary File 2. The mass spectrometry proteomics data have been deposited to the ProteomeXchange Consortium via the PRIDE^58^ partner repository with the dataset identifier PXD052173.

### RNA extraction and reverse-transcription quantitative PCR (RT-qPCR)

Total RNA was extracted from cells using RNAiso Plus (Takara, # 9109) according to the manufacturer’s instructions. cDNA was synthesized using the ABScript II cDNA first strand synthesis kit (ABC clone, # RK20400). qPCR reaction was performed by using SYBR Green qPCR Master Mix (TargetMol, # C0006), β-actin was used as the internal reference gene. All primer sequences were listed in Supplementary Table 1.

### Cell proliferation and viability assay

1× 10^3^ PC3 or DU145 cells, 2× 10^3^ 22Rv1 cells per well were seeded in 96-cell plates for cell proliferation. 2× 10^3^ DU145 cells or 2× 10^4^ 22Rv1 cells per well were seeded in 96-cell plates for cell viability test under different drug stimulations. 10μL/well CCK-8 regent (CCK-8, TargetMol, # C0005) was added for 4 hours before the indicated time points, and then the data were collected by measuring the absorbance at 450 nm. Values were obtained from triplicate wells.

### Trans-well assay

Cells were trypsinized and rinsed twice with PBS, followed by resuspended in serum-free medium at concentrations of 2 × 10^5^ for 22Rv1, 3 × 10^5^ cells / mL for DU145 and PC3 cell lines. A 200 μL aliquot of the single-cell suspension was carefully pipetted into Trans-well inserts, either with or without VBIT4 (TargetMol, # T13287). Concurrently, the lower chambers were filled with 750 μL complete culture medium containing 10% FBS. After incubating for the indicated durations, the inserts were rinsed twice with cold fresh PBS, then fixed with 3.7% formaldehyde solution for 5 min at room temperature, followed by another twice rinse. Permeabilization was achieved using cold methanol at −20 ℃ for 20min, again followed by double rinse. The cells were ultimately stained with 0.1% Crystal Violet Aqueous Solution (Solarbio, # G1064) for 30 min at room temperature, after which excess dye was wiped off with a cotton swab. The migrated cells were observed and photographed using an Inverted Microscope (Leica, DMi8 automated) or Stereo Microscope (Carl Zeiss Microscopy GmbH). The migration rate, in terms of number or area, was quantitatively analyzed using ImageJ software.

### Colony formation assay

500 cells for DU145 or PC3 cells, 1000 cells for 22Rv1 per well were seeded on 24-well plates for 24 hours before relevant treatment. The medium was changed every three days. At the end time point, cells were stained with 0.1% Crystal Violet Aqueous Solution. The colony (with >50 cells) number was counted by ImageJ software.

### Wound healing assay

Cells were seeded onto 24-well plates at a density of 2×10^5^ for DU145 or PC3 cells and 2×10^6^ for 22Rv1 cells until confluence. And the plates were covered with three evenly and straight location lines across the back by a mark-pen ahead of the scratch. A 200μL micropipette tip was used to create three evenly distributed scratches, perpendicular to the location line, then washing the wells three times with PBS to remove the floating cells. Then, the complete medium with or without VBIT4 was added into the well. The plates were observed and photographed using an inverted microscope (Leica, DMi8 automated) initially at 0 h and subsequently at the designated endpoint. The wound area was calculated using ImageJ software. The relative migration rate was calculated using the following formula: (wound area at 0 h - wound area at endpoint) / wound area at 0h × 100%. Three replicates were set for each experiment, and a minimum of three paired pictures were collected per replicate to ensure reproducibility and reliability of the data.

### Transmission Electron Microscope (TEM)

For the analysis of mitochondrial ultrastructure, 22Rv1 cells were fixed with 2.5% glutaraldehyde (resolved in 10 mM PBS (pH 7.0-7.5)) for 2 to 4 hours at 4 ℃. Then the cells were pre-embedded in 1% agarose and post-fixed with 2% ferrocyanide-reduced osmium tetroxide for 2 h at room temperature in the dark, followed by washing three times with PBS (100 mM, pH 7.4). Subsequently, the dehydrated cells were embedded in resin and sectioned to a thickness of 60-80 nm. The sections were stained with 2% uranium acetate saturated alcohol solution in the dark for 8 min, followed by three washes in 70% alcohol and ultra-water, respectively. Additionally, the sections were stained with 2.6% lead citrate solution for 8 min in a CO_2_-free environment. The samples were then photographed using a transmission electron microscope.

### Mitochondria analysis

Mito-Tracker Deep Red (Beyotime, # C1032) was used to stain the mitochondria, and the nuclei was stained with Hoechst 33342 (TargetMol, # T5840). Briefly, cells were seeded onto the laser confocal dishes for 48 hours. The cells were washed twice with warm, serum-free medium and covered with 200 nM Mito-Tracker Deep Red solution (diluted with warm, serum-free medium), then the cells were incubated at 37 ℃ for 30 minutes in the dark. The labelled cells were washed twice as above, and covered with 3 μg/mL Hoechst solution at 37 ℃ for 15 minutes in the incubator. After another two times washing procedure, 1mL warm, complete culture medium with or without VBIT4 was added onto the cells. A high-resolution laser confocal microscope (Leica) with 63x/1.40 Oil DIC objective was used to observe and the labeled cells were photographed, which were lied at 37 ℃ with humidified atmosphere of 5% CO_2_.

### Quantification of Mitochondrial membrane potential (MMP, Δψm)

Mitochondrial membrane potential was assessed using JC-1 probe (Solarbio, # M8650), as previously described with modifications^59^. 2×10^5^ DU145 cells were seeded onto 24-cell plates overnight and treated with or without 5μM VBIT4 for 24 hours. After discarding the culture medium, the adherent cells were washed twice with fresh PBS. Subsequently, 300μL of JC-1 working solution was added into the well. The cells were then incubated in the dark at 37 ℃ for 20 minutes. The labeled cells were washed twice with the cold washing buffer, then trypsinized and resuspended in PBS with 2% FBS for quantification using a flow cytometer (Beckman). JC-1 monomers and aggregates were detected at PE and FITC channels, respectively. The positive control treated with 10mM CCCP for 24 hours was used to adjust compensation, detailed in Supplementary figure 7. 30000 cells were collected per sample.

### Cell death assay

Propidium iodide (Beyotime, # ST511) staining was performed to assess cell death, as previously described ^8, 60^. Briefly, 2×10^5^ cells were seeded on 24-cell plates for 24 hours before treatment. After relevant stimuli, cells were collected (including floating dead cells) and washed twice with PBS (Solarbio, # P1010), then incubated with 500μL PI staining solution (3μg/ml) in PBS for 30min at 37 ℃. Next, cells were resuspended in 200μL PBS with 2% FBS, and the cell death rate was measured using flow cytometer. At least, 1×10^4^ single cells were collected per well, and three replicates were set for each experiment.

### Quantification of Lipid Peroxidation

To quantify the level of lipid peroxidation of various treatment cells, BODIPY™ 581/591 C11 (Thermo Scientific, # D3861) was used as previously described^7, 9^. 2×10^5^ cells were seeded onto 24-well plates overnight, and treated with or without RSL3 (TargetMol, # T3646), Ferrostatin-1 (TargetMol, # T6500) or VBIT4 (TargetMol, # T13287) for 24 hours. Cells were then washed twice with PBS and incubated with culture medium (without FBS and PS) containing 5μM BODIPY™ 581/591 C11 at 37 ℃ for 30min in dark. Labeled cells were washed twice with PBS to remove the excess dye, and finally resuspended in cold PBS containing 2% FBS for flow cytometer analysis. A minimum 10000 events were collected to detect the oxidized C11-BODIPY at FITC channel.

### Quantification of Mitochondria ROS

As previously reported, MitoSOX (MCE, # HY-D1055) was used to assess the level of mitochondrial superoxide in the study^7^. Briefly, 2×10^5^ cells were seeded on 24-well plates overnight, then treated with or without VBIT4 (1μM for 22Rv1, 5μM for DU145) for 24 hours. The cells were washed twice with fresh PBS and stained in serum-free medium containing MitoSOX at 37 ℃ for 30 min in dark. The labeled cells were washed and resuspended in PBS, then detected using flow cytometry at PE channel. All flow cytometry experiments were performed at least in triplicate, and the data was processed with FlowJo (version 10.8).

### Quantification of Mitochondria Lipid Peroxidation

MitoPeDPP (Dojindo, # M466) was employed to quantify the level of mitochondria lipid peroxidation in prostate cancer cells, according to the manufacture’s protocols. Briefly, 5× 10^4^ DU145 cells were seeded onto a 35mm laser confocal dish (Biosharp) and cultured for 48 h. The cells were washed twice with warm, serum-free medium to remove any debris. Subsequently, the cells were cultured with 0.5 μM MitoPeDPP in conjunction with 200 nM Mito-Tracker Deep Red solution (diluted with warm and serum-free medium) in the dark at 37 ℃ for 30 min. After that, the cells underwent another two rinses with the same medium. Then 1mL warm and complete culture medium with or without VBIT4 was added to the cells. Finally, the labeled cells were imaged using a high-resolution laser confocal microscope (Leica) with 63x/1.40 Oil DIC objective, at 37 ℃ with humidified atmosphere of 5% CO_2_.

### Immunohistochemistry (IHC) and Hematoxylin-Eosin staining (H & E)

A prostate cancer tissue microarray (Lot No. PRC1601) containing 78 paired tumor and adjacent tissues was obtained from Outdo Biotech (Shanghai, China) to analyze the expression levels of VSTM2L. After the standard procedure of immunohistochemistry of our laboratory^57^, the staining index of VSTM2L was calculated as the following formula: the staining intensity × the scope of stained-positive cells. For staining intensity, the score was evaluated on a 4-point scale: 0 (no appreciable staining, negative), 1 (faint yellow, weak intensity), 2 (yellow-brown, moderate intensity), 3 (brown, strong intensity). The scope of the stained-positive cells was divided into four categories: 1 (< 25% positive cells), 2 (26–50% positive cells), 3 (51–75% positive cells), and 4 (>75% positive cells). Finally, the expression of VSTM2L was divided into high and low levels according to their mean scores.

For the analysis of immunohistochemistry and H & E staining for xenograft, the fixed tumor tissues were paraffin-embedded subjected to the standard procedures. The primary antibodies utilized were anti-Ki67, anti-VSTM2L and anti-GPX4. The images were captured by an upright microscope (Zeiss).

### VDAC1 cross-linking assay

The VDAC1 cross-linking assay was performed as previously reported with modifications^24, 45^. Briefly, cells were harvested and washed twice with cold PBS (pH 8.0), followed by centrifuging at 3000g for 5 min at 4 ℃. The collected pellets were resuspended in 740 μL of 0.5mM Ethylene glycol-bis (succinic acid N-hydroxysuccinimide ester) (EGS, Thermo, # 21565) solution diluted with cold PBS (pH 8.0), and incubated for 30min at room temperature. To quench the reaction, 10 μL of 1.5 M Tris-HCL (pH 7.8) was added, and the reaction mixture was cultured at room temperature for another 15 min. Finally, the cultured cells were harvested in lysis buffer and the protein concentration was measured using BCA assay. 50 μg of the protein samples were subjected to 7.5% SDS-PAGE and immunoblotting using an anti-VDAC1 antibody.

### Measurement of Glutathione (GSH)

For the purpose of assessing glutathione (GSH) concentrations in cellular and tissue samples, we utilized the Glutathione Assay Kit (Nanjing Jiancheng, # A006-2), following the manufacturer’s guidelines. All experiments were normalized based on the total protein concentration and were performed in triplicate.

#### Immunofluorescence (IF) assay

5 × 10^4^ DU145 and PC3 or 1 × 10^5^ 22Rv1 cells per well were seeded onto 14 mm glass slide in 24-well plates and cultured for at least 24 h. The cells were washed twice with 1 × PBS to remove any debris. Subsequently, the cells were cultured with 200 nM Mito-Tracker Deep Red solution (diluted with serum-free medium) in the dark at 37 ℃ for 30 min. After that, the cells underwent another two rinses with 1 × PBS. Then the cells were fixed with cold methanol in the dark at 4 ℃ for 15 min. Another 3 times washing with 1 × PBS was following. The fixed cells were blocked with 5% BSA for 1 hour and incubated with ideal primary antibodies (anti-HK2, anti-VSTM2L and anti-VDAC1) in the dark at 4 ℃ overnight. The cells were washed 3 times with cold 1 × PBS and incubated with the fluorescent secondary antibody (diluted with 5% BSA) in the dark at room-tempreature for 1 hour. Followed another 3 times washing with 1 × PBS and the Hoechst (5 μg / mL) was used to stain the cell nucleus. Finally, the stained cells were imaged using a high-resolution laser confocal microscope (Leica) with 63x/1.40 Oil DIC objective.

#### Mitochondria and Cytoplasmic proteins extraction assay

In order to separately extract the mitochondria and cytoplasmic proteins from prostate cancer cell lines, a commercial Cell Mitochondria Isolation Kit (Byotime, # C3601) was used according to the manufacture. All experiments were performed in triplicate.

## Statistical analysis

All experiments were conducted at least in triplicate. GraphPad Prism 9 was used to obtain statistical *p*-values by application of student’s *t*-test or ANOVA for group comparison. Kaplan-Meier method was used to plot the PFS and DFS curves, compared by log-rank test. FlowJo (version 10.8) was employed to quantify the data acquired using flow cytometer. ImageJ software was utilized to analyze the morphological parameters of mitochondria. Data were presented as mean ± standard deviation (SD). For all tests, *p* < 0.05 was considered significant (ns: not significant).

## Data availability

The VSTM2L gene expression in Prostate adenocarcinoma (PRAD) was derived from UALCAN (https://ualcan.path.uab.edu/analysis.html), GEPIA (http://gepia.cancer-pku.cn/index.html) and GENT2 (http://gent2.appex.kr/gent2/) databases. The survival curves of VSTM2L were derived from cBioPortal (https://www.cbioportal.org/). All data underpinning the conclusions of this research are accessible through a reasonable request made to the corresponding author.

## Acknowledgements

This work was funded by the Natural Science Foundation of Shaanxi Province (2023-JC-YB-716), Fundamental Research Funds for the Central Universities (GK202201004), Excellent Graduate Training Program of Shaanxi Normal University (LHRCTS23091).

## Author contributions

P.G., and X.M.D. conceived and designed the project. J.Y., X.L. assistant by J.L.H., L.L., Y.T.R., and X.N.A performed the experiments. J.Y. and X.M.D performed data analysis. P.G., X.M.D., and J.Y. wrote the manuscript.

---

**Supplementary figure 1.** Representative image of an SDS-PAGE gel stained with Coomassie blue. LNCaP cells transfected with Flag-VDAC1 were collected for immunoprecipitation with anti-Flag antibodies, and separated by SDS-PAGE gel. The gel was subsequently stained with Coomassie blue to visualize the distinct binding bands. The source data was provided as supplementary table 3 and supplementary table 4.

**Supplementary figure 2. VSTM2L is a novel oncogene in PCa. A** The mRNA levels of *VSTM2L* in PRAD and normal tissues from UALCAN database. **B** The mRNA levels of *VSTM2L* in the indicated cell lines. Data shown as mean ± SD, n=3. **C-E** Association between Disease-free survival of prostate cancer patients and *VSTM2L* mRNA levels from the cBioPortal database. **F, G** RT-qPCR and western blot assays were performed to detect the mRNA (**F**) and protein (**G**) levels of VSTM2L, respectively, in PCa cells transfected with VSTM2L shRNAs or control shRNA. Data shown as mean ± SD, n=3. **H, I** Proliferation (**H**) and colony formation (**I**) abilities in PC3 cells with downregulation of VSTM2L. Data shown as mean ± SD, n=3. **J-M** Migration ability of VSTM2L knockdown PC3 cells was measured by wound healing (**J, L**) and trans-well (**K, M**) assay, respectively. Data shown as mean ± SD, n=3, Scale bars = 250 μm. *P* value was determined by log-rank test (**C, D, E**) and one-way (**B, F, I, L, M**) or two-way (**H**) ANOVA.

**Supplementary figure 3. Supression of VSTM2L changes the morphology of PCa cells and mitochondria. A** Representative images of transmission electron microscopy (TEM) for 22Rv1cells transfected with VSTM2L shRNA or control shRNA. Scale bars = 5 μm. **B** Representative images of high-resolution laser confocal microscopy for the morphology of mitochondria in 22Rv1 cells transfected with shVSTM2L or control. Scale bars = 5 μm. **C** Mitochondria network for images shown in **Fig. 3 C** were determined by ImageJ. Data shown as mean ± SD of n=3 technical replicates. *P* value was determined by unpaired two-tailed Student’s *t*-test. **D** The DU145 cells with VSTM2L shRNA (shVSTM2L-1, shVSTM2L-2) or control shRNA (Control) were stained with Annexin V and analyzed by flow cytometry. Data shown as mean ± SD of n=3 technical replicates. **E** Western blotting analysis of the protein levels of apoptosis markers (Bcl-2 and Bax) of DU145 cells transfected with VSTM2L shRNA or control shRNA. Protein levels were normalized to total GAPDH. **F** Representative and Quantification of flow cytometry histograms show the percentage of cell death in DU145 cells transfected with VSTM2L shRNAs or control shRNA, stained by PI (3 μg/mL). Cells were treated with or without Fer-1 (5 μM) for 24h. Data shown as mean ± SD, n=3, ns not significant. *P* value was determined by one-way ANOVA (D) and two-way ANOVA (F).

**Supplementary figure 4. VSTM2L overexpression inhibited ferroptosis of PCa cells. A** Western blot analysis of the VSTM2L protein levels in VSTM2L stably overexpressed PCa cells and control cells. GAPDH was used as a loading control. **B** RT-qPCR analysis of the *VSTM2L* mRNA levels in VSTM2L stably overexpressed prostate cancer cells and control cells. Data shown as mean ± SD, n=3. **C, D** Cell death was detected by PI staining (3μg/mL) and quantified by flow cytometer in DU145 cells with overexpression of VSTM2L. Data shown as mean ± SD, n=3. **E** Western blot analysis of GPX4 and VSTM2L protein levels in VSTM2L overexpressed or control cells. **F** The level of glutathione (GSH) in VSTM2L overexpressed or control cells. Data shown as mean ± SD of n=3 technical replicates. *P* value was determined by unpaired two-tailed Student’s *t*-test.

**Supplementary figure 5. VSTM2L regulates mitochondria ROS level in PCa cells. A** The mtROS levels of PCa cells with VSTM2L overexpression were evaluated by a MitoSOX (5 μM) probe staining. Data shown as mean ± SD, n=3. **B, C** VSTM2L suppressed 22Rv1 cells cultured with or without VBIT4 (1 μM) for 24h were stained by a MitoSOX (5 μM) probe and analyzed by flow cytometry to evaluate the mtROS levels. Data shown as mean ± SD, n=3. **D** The mtROS levels of Pca cells with VSTM2L suppression were treated with or without MitoQ (10 nM) for 24h, then stained by a MitoSOX (5 μM) probe and analyzed by flow cytometry to check mtROS levels. Data shown as mean ± SD, n=3. *P* value was determined by unpaired two-tailed Student’s *t*-test (**A**) and two-way ANOVA (**C, D**).

**Supplementary figure 6. Representative images of high-resolution laser confocal microscopy for the morphology of mitochondria in VSTM2L knockdown DU145 cells or control cells cultured with or without VBIT4.** The mitochondria was stained with Mitotracker (200 nM), and nucleus with Hoechst (3 μg/mL). Scale bars = 50 μm.

**Supplementary figure 7. Gating strategies of flow cytometer. A** Gating strategies for cell death, lipid peroxidation and mtROS by flow cytometer. **B** Gating strategy for JC-1 staining by flow cytometer.

