## Supplementary figures and images for "VSTM2L protects prostate cancer cells against ferroptosis via inhibiting VDAC1 oligomerization and maintaining mitochondria homeostasis"

### Supplementary Figure 1

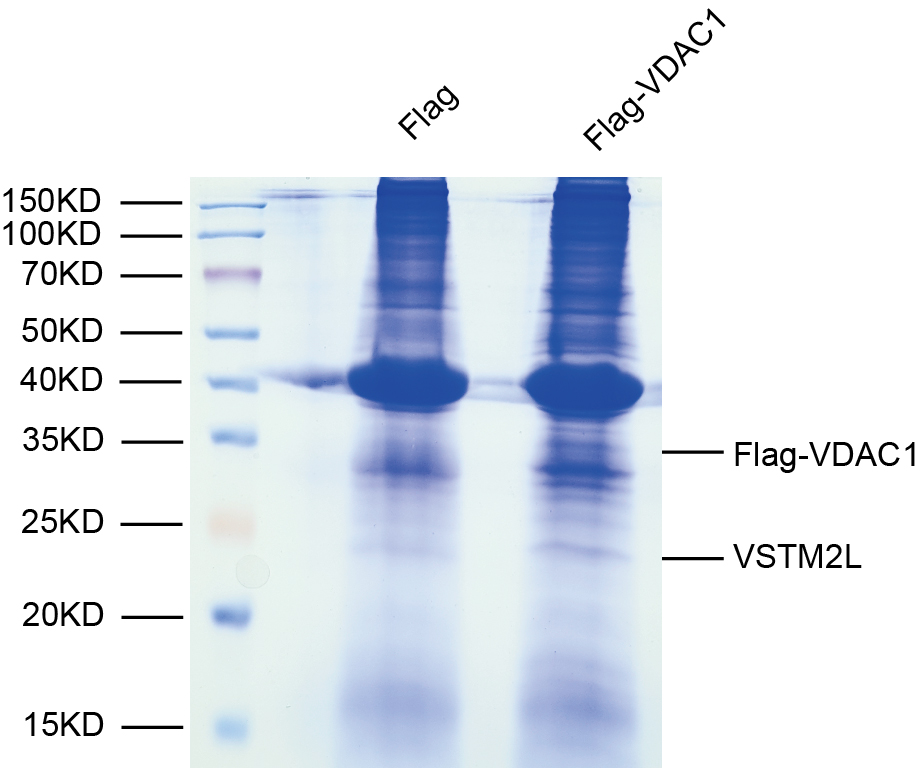

### Supplementary Figure 2

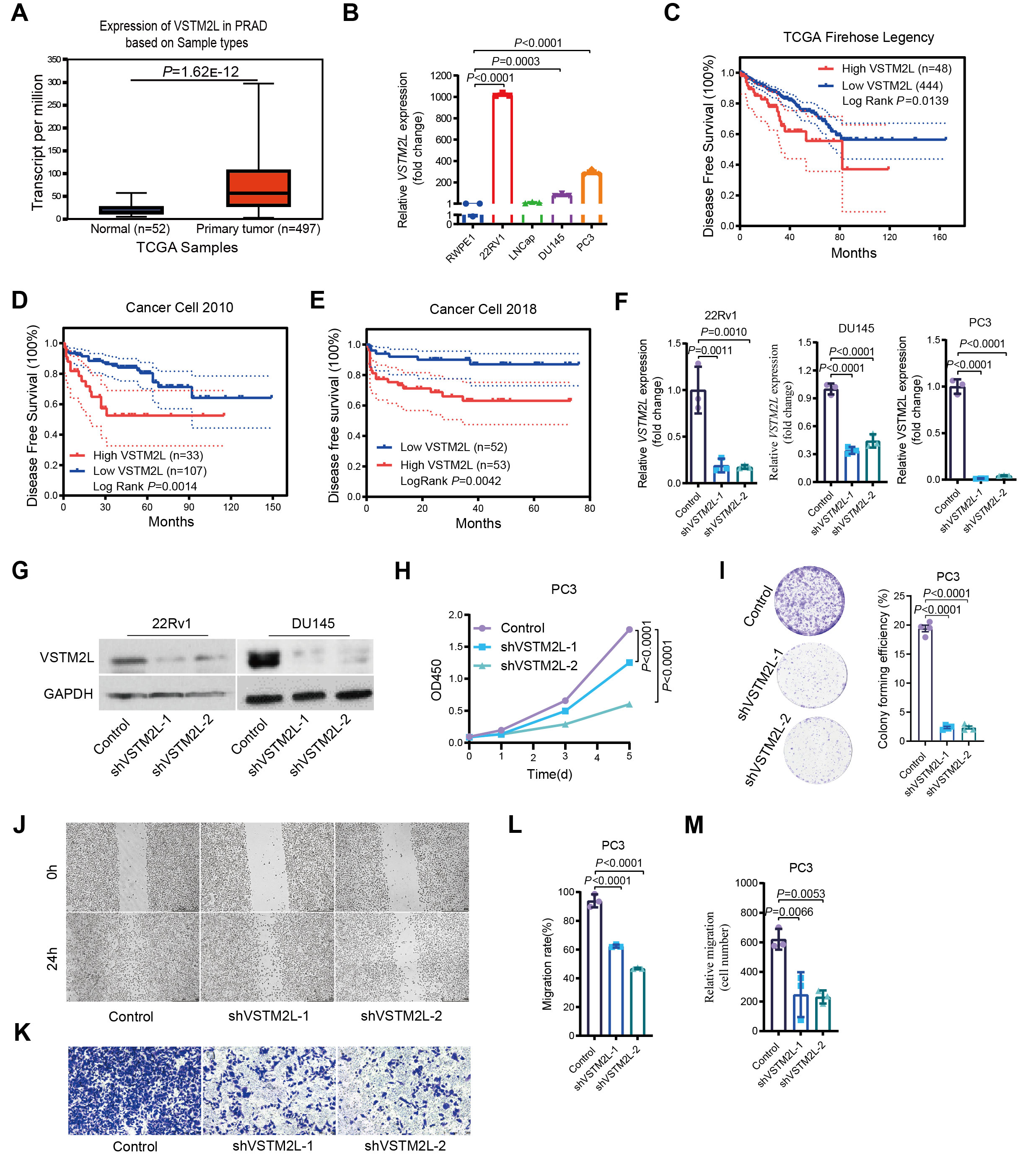

### Supplementary Figure 3

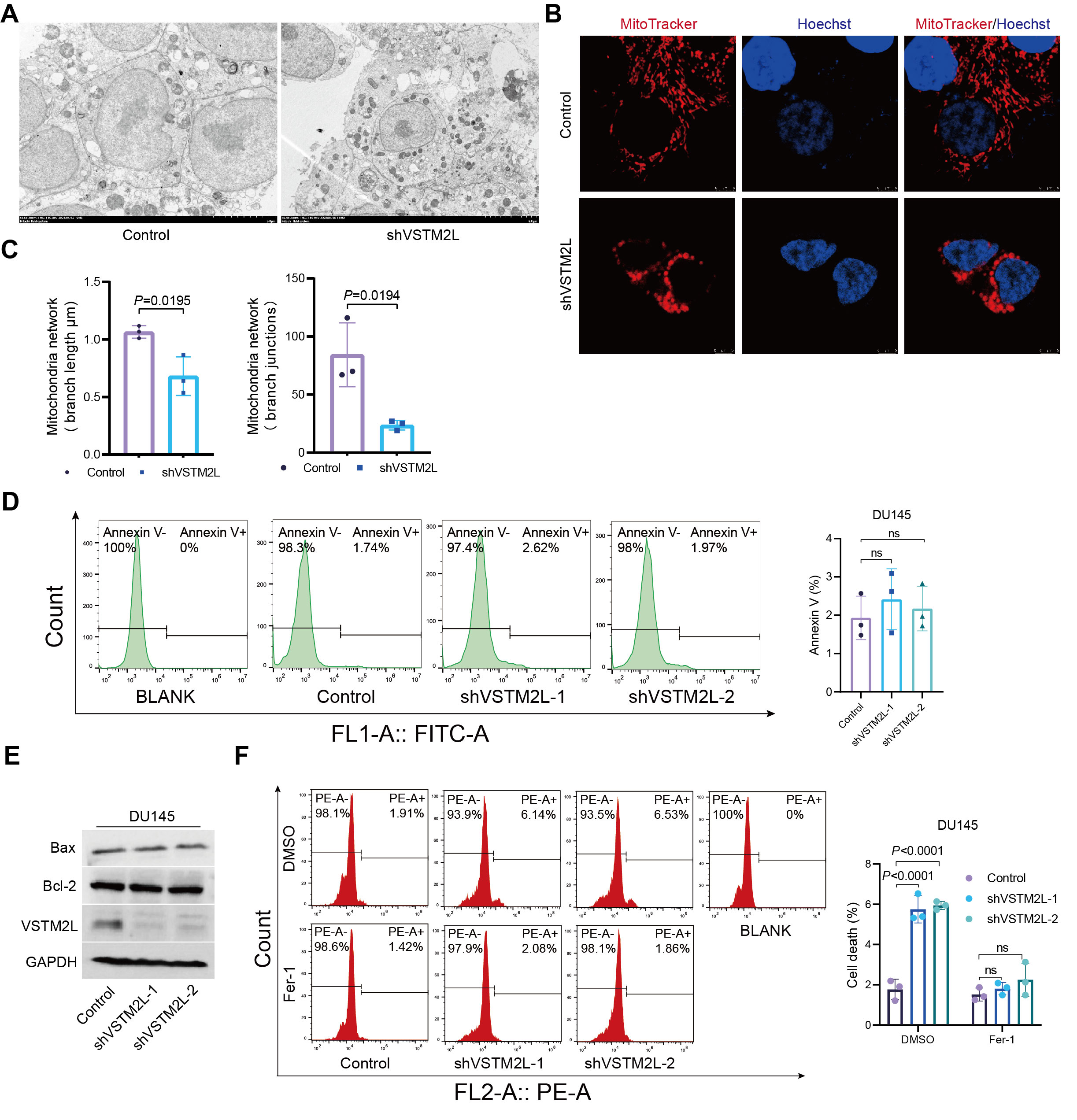

### Supplementary Figure 4

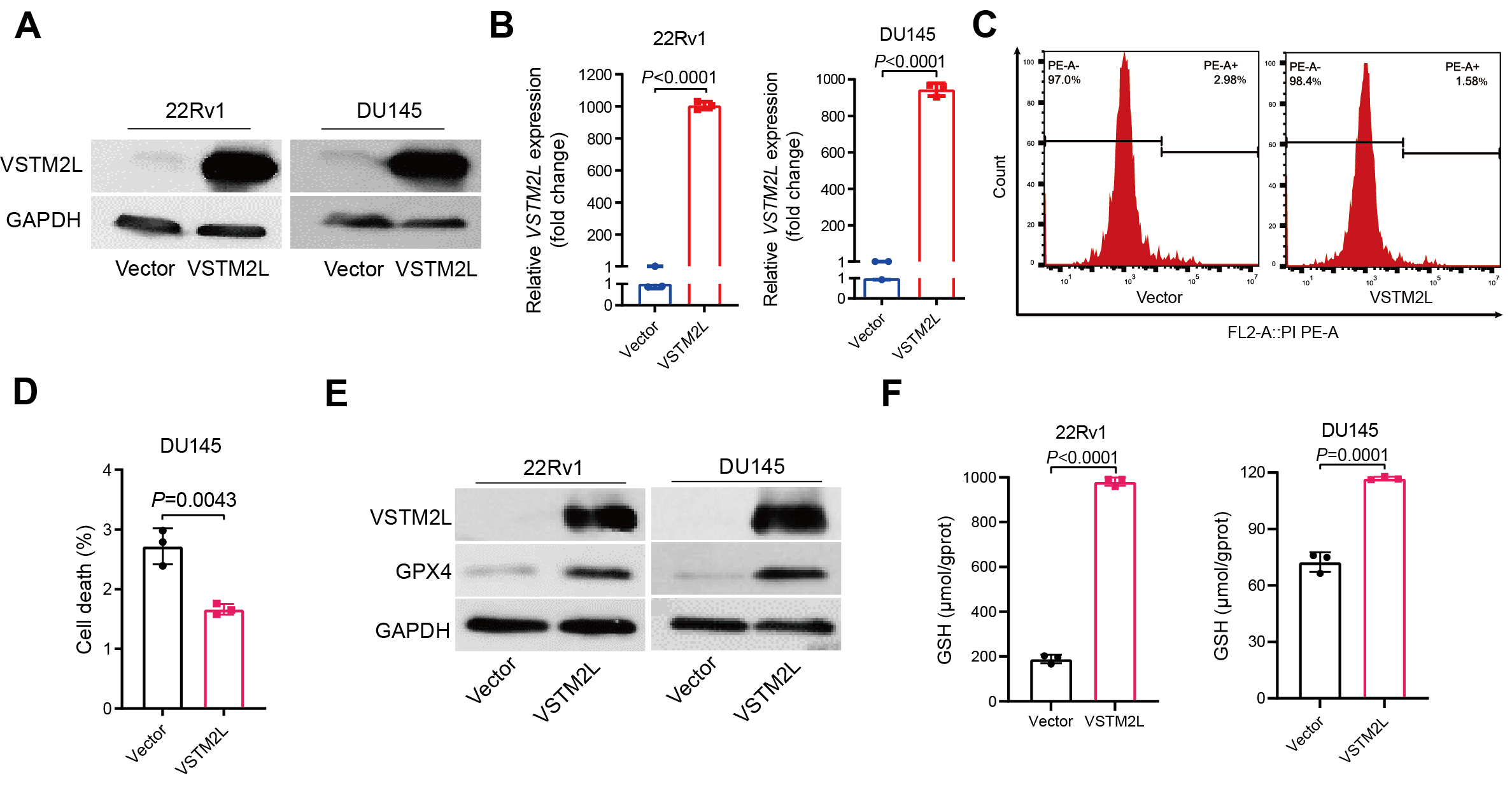

### Supplementary Figure 5

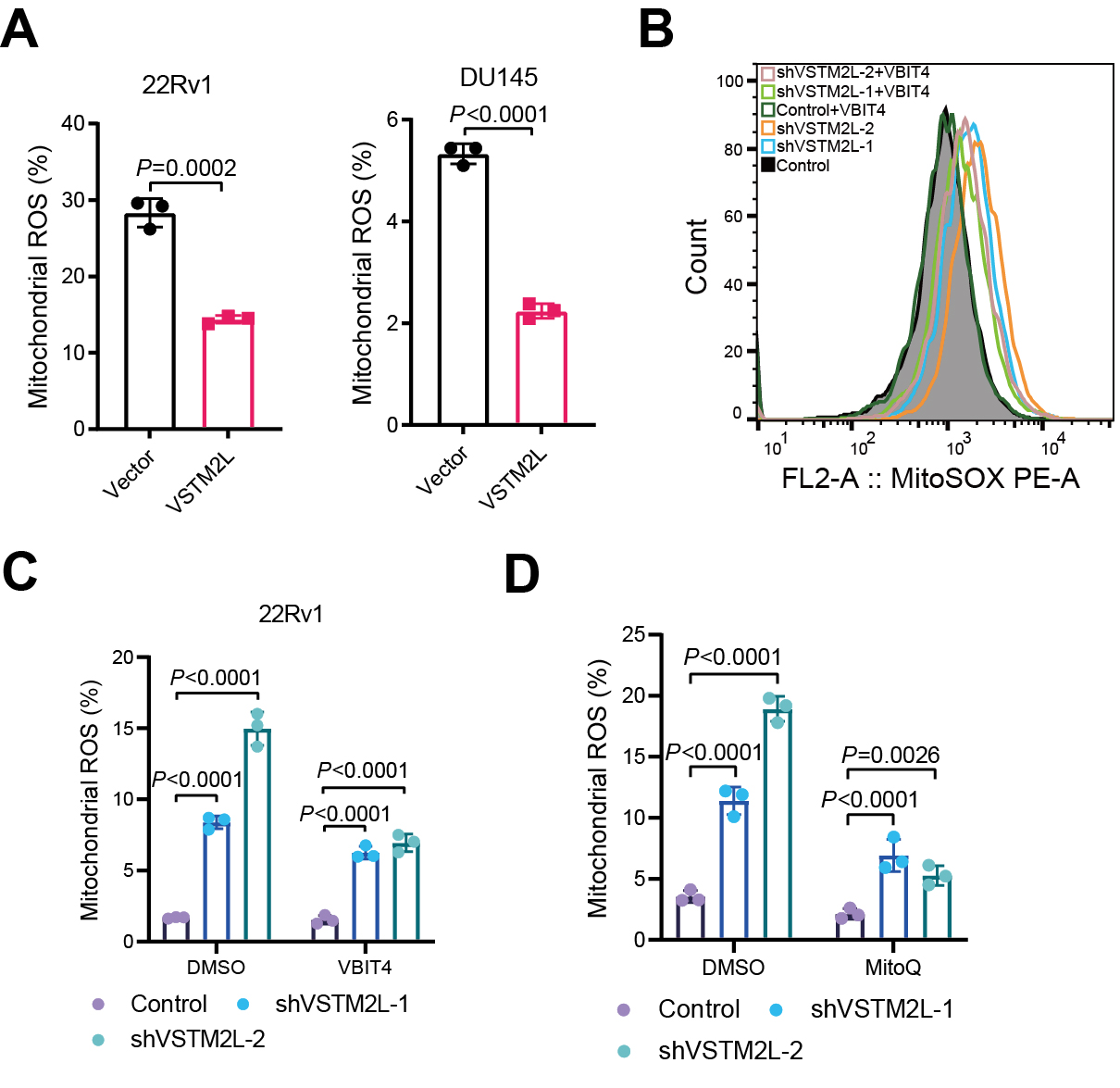

### Supplementary Figure 6

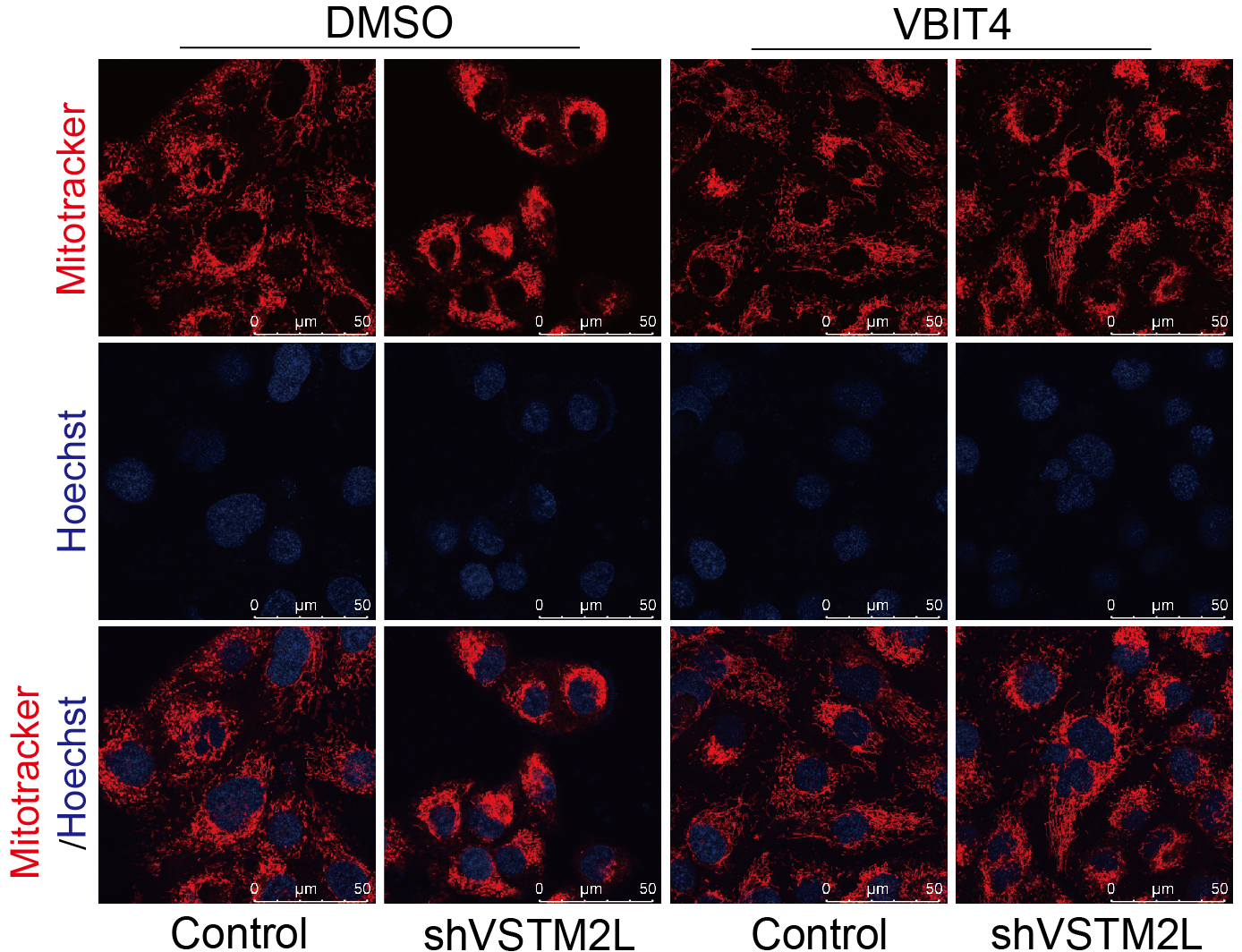

### Supplementary Figure 7

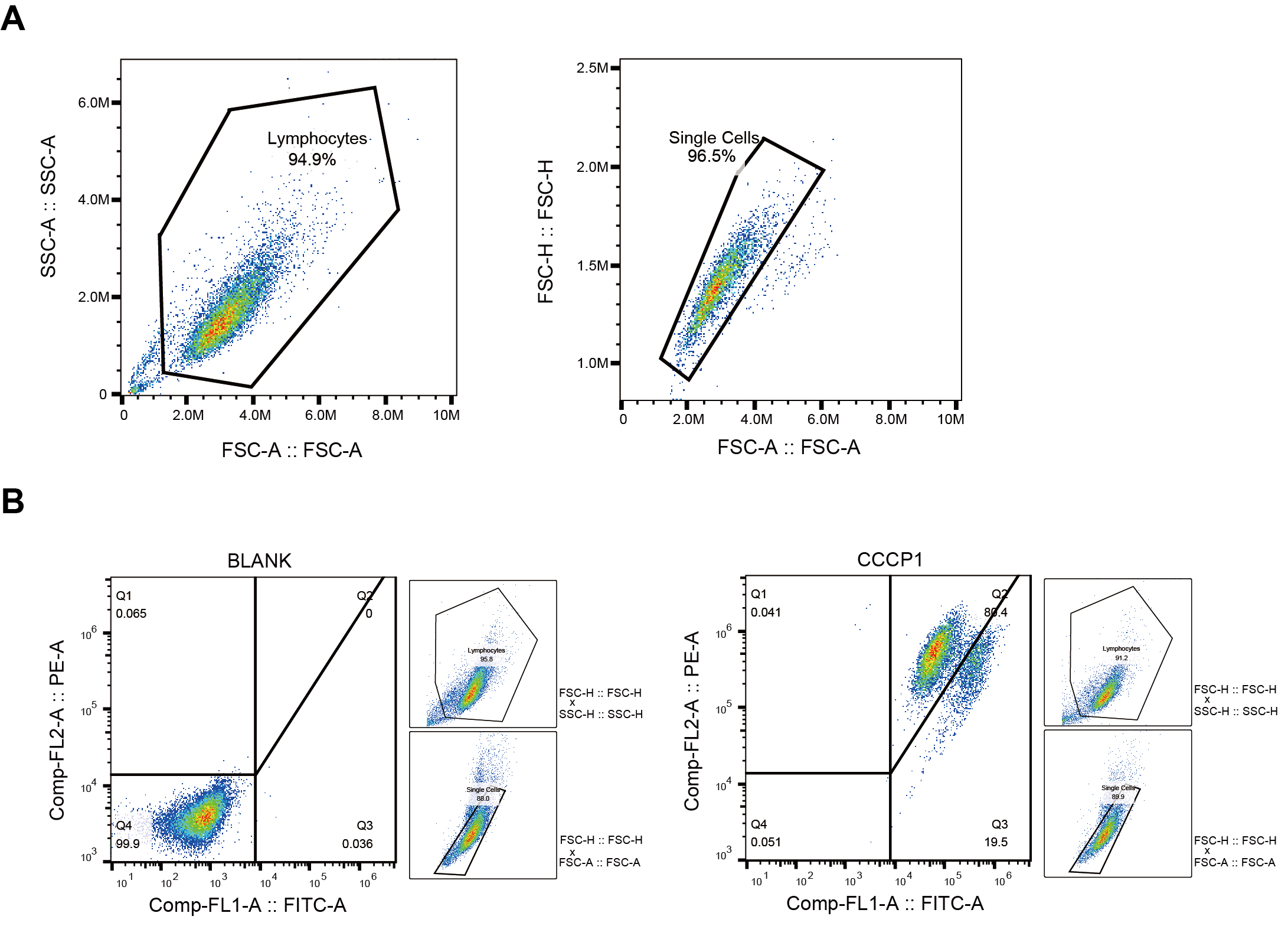
